# *e*^−2^ Threshold for Bistability in Non-equilibrium Sorting of Biomolecular Condensates

**DOI:** 10.64898/2026.09.28.754880

**Authors:** Weifeng Chen, Shiyuan Pei, Guozhi Yu

**Affiliations:** College of Life Sciences, Sichuan Agricultural University, Ya’an 625014, China

## Abstract

Cells use ATP-driven chaperones to extract clients from biomolecular condensates selectively, yet no closed-form criterion links molecular parameters to condensate quality control. We derive a minimal non-equilibrium sorting reactor model that reduces the reaction-diffusion dynamics to one algebraic equation. In the idealized low-load limit, a concentration-dependent partition coefficient produces a saddle-node bifurcation between functional clearance and pathological storage. The critical Damköhler number is Da_0*c*_ = *e*^*−*2^ and is independent of the feedback strength. Corrections shift the value but preserve the topology. The model predicts a Kramers-type functional lifetime and recovery costs that scale as *R*^3^. Critical slowing down near the bifurcation provides early-warning signatures for FCS and FRAP.

## Introduction

*—*Biomolecular condensates formed by liquid-liquid phase separation (LLPS) enrich client proteins and nucleic acids selectively [1–3]. The multivalent interactions that drive this selectivity also make condensates susceptible to pathological liquid-to-solid transitions, as seen for FUS and TDP-43 in ALS and related diseases [4–7]. ATP-consuming chaperones counter this by extracting misfolded clients from condensates such as nucleoli and stress granules. Quantitative imaging and biochemical experiments show that chaperones discriminate between clients with similar equilibrium affinities [8– 12]. Reviews have highlighted the crosstalk between condensates and proteostasis [13–15], and chaperones are now recognized as regulators of condensate formation and dispersal [16–18]. Condensate quality control is therefore a kinetically controlled, non-equilibrium process. The central question is what sets the threshold between functional clearance and pathological storage.

Active-droplet and active Cahn-Hilliard theories have shown that chemical reactions can control droplet size and suppress or accelerate Ostwald ripening [19–26]. These approaches rely on numerical integration. They do not give closed-form relations that link molecular parameters to fidelity, stability, and energy consumption. Equilibrium Flory-Huggins theory cannot describe the non-equilibrium sorting. The dual role of ATP, which activates chaperones and acts as a hydrotrope, has not been incorporated into a threshold criterion [27–32].

We develop here a minimal non-equilibrium sorting reactor (MNSR) model. It reduces the reaction-diffusion dynamics of active client extraction to one algebraic equation. The reduction reveals a saddle-node bifurcation between functional clearance and pathological storage, and in the idealized low-load limit yields a closed-form threshold. Molecular corrections shift the threshold but preserve the topology. The same bifurcation predicts critical slowing down near the threshold.

### The MNSR model

Consider a spherical condensate of radius *R* surrounded by a large reservoir. The droplet is rich in scaffold and recruits client molecules selectively from the dilute phase. Two fluxes compete. Clients enter by passive diffusion across the interface, driven by the concentration mismatch between outside and inside. Inside the droplet, ATP-fueled chaperones extract clients actively and return them to the reservoir.

When diffusion inside the droplet is fast and the external concentration field adapts quickly, the spatial degrees of freedom can be eliminated. The system reduces to a well-mixed reactor in which the intra-droplet concentration is controlled by the balance of supply and removal. See Supplemental Material at [URL] for this coarse-graining from the full Flory-Huggins free energy with active reactions. For a mixture of *n* client species, the steady-state condition takes the dimensionless form

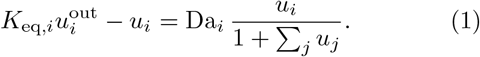

Here 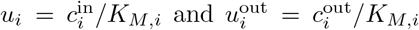 are the dimensionless intra-droplet and external concentrations, with *K*_*M,i*_ the chaperone Michaelis constant for client *i*. The mass-transfer coefficient is 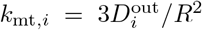. The Damköhler number Da_*i*_ ≡ *K*_eq,*i*_*k*_ext,*i*_*/k*_mt,*i*_ compares active extraction to diffusive supply. The denominator 1 + ∑_*j*_ *u*_*j*_ captures competition among clients for the shared chaperone pool. In the single-client, low-load limit considered below, we replace *K*_eq_ by its zero-concentration baseline *K*^0^, which gives the constant clearance strength

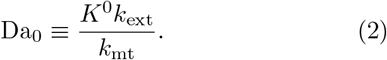

Equation (1) is the general MNSR equation.

Now focus on a single client species and suppose its intra-droplet concentration is far below the chaperone saturation threshold, *u* ≪ 1, where *u* = *c*^in^*/K*_*M*_. The denominator simplifies to unity and extraction follows linear kinetics. The steady-state balance, already in dimensionless form, reads

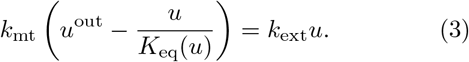

Here *k*_mt_ is the mass-transfer coefficient defined above, *k*_ext_ is the linear extraction rate, and *K*_eq_(*u*) emphasizes that the partition coefficient may depend on the intra-droplet concentration. Equation (3) is the single-client MNSR equation with an arbitrary concentration-dependent partition coefficient.

The partition coefficient *K*_eq_(*u*) now depends on the intra-droplet concentration. If client molecules do not interact with one another, *K*_eq_ is a constant set by the scaffold-client affinity. Many clients that enter condensates form weak homotypic contacts. Each incoming molecule displaces solvent and lowers the free-energy cost for the next entry. Supplemental Material Sec. S2.A shows that these interactions add a term linear in *u* to the exchange chemical potential. Equating the chemical potentials across the interface gives the partition coefficient an exponential concentration dependence,

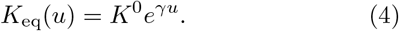

The constant *K*^0^ is the baseline affinity at infinite dilution. The dimensionless parameter *γ* is positive only when direct client-client attractions overcome the entropic penalty of excluded volume, i.e., 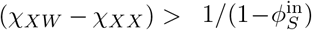, as shown in Supplemental Material Sec. S2.A. It condenses two physical effects. Excluded-volume entropy contributes because each client that enters the dense phase displaces solvent, and direct attractive interactions among clients provide an additional enthalpic drive. As the client concentration rises, the condensate becomes more receptive to further entry.

With this exponential feedback the steady-state condition takes a compact form. Substituting Eq. (4) into Eq. (3) and multiplying by *K*^0^*/k*_mt_, we define the dimensionless external supply

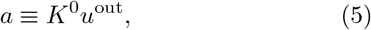

so that the steady-state condition becomes

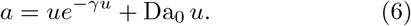

The external supply *a* is the dimensionless intra-droplet concentration that would equilibrate in the absence of feedback and extraction. The clearance strength Da_0_ weighs active removal against passive resupply. Equation (6) is the equation of state for the condensate. Its roots determine whether the droplet operates in a functional clearance mode or collapses into a pathological storage mode. The fate of the condensate depends on three dimensionless parameters, *γ, a*, and Da_0_.

### The threshold of bistability

*—*Equation (6) determines the steady states. Its right-hand side defines Φ(*u*) = *ue*^*−γu*^ + Da_0_ *u*, whose derivative Φ′(*u*) = (1 − *γu*)*e*^*−γu*^ + Da_0_ controls the number and stability of solutions. When Da_0_ is large, Φ′(*u*) *>* 0 everywhere and a single stable clearance state exists. For Da_0_ *< e*^*−*2^, however, Φ(*u*) develops a local maximum and minimum, giving three fixed points. They are a stable clearance state *u*_low_, an unstable saddle *u*_barr_, and a stable storage state *u*_high_. This bistable structure is visualized in the phase portraits of Fig. 2(a). At low supply only the clearance attractor is present, while inside the bistable window two stable states coexist separated by the saddle. The storage state, marked by a red border in Fig. 2(a), corresponds to runaway accumulation. Crossing the upper saddle-node boundary *a* = *a*_max_ eliminates the clearance state and leaves only the pathological attractor. The corresponding bifurcation diagram in Fig. 2(b) traces these fixed points as *a* varies, with stable branches solid, the saddle dashed, and the bistable window shaded pink between *a*_min_ and *a*_max_. The bistable region in the (Da_0_, *a*) plane is bounded by the two saddle-node lines, which meet at the cusp

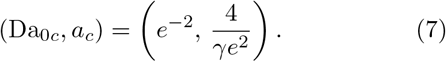

Within the exponential-feedback model this threshold is parameter-free. The critical clearance strength Da_0*c*_ = *e*^*−*2^ does not depend on *γ*, while the critical supply *a*_*c*_ = 4*/*(*γe*^2^) scales as 1*/γ*. The feedback strength *γ* only rescales the concentration and supply axes.

**FIG. 1.**
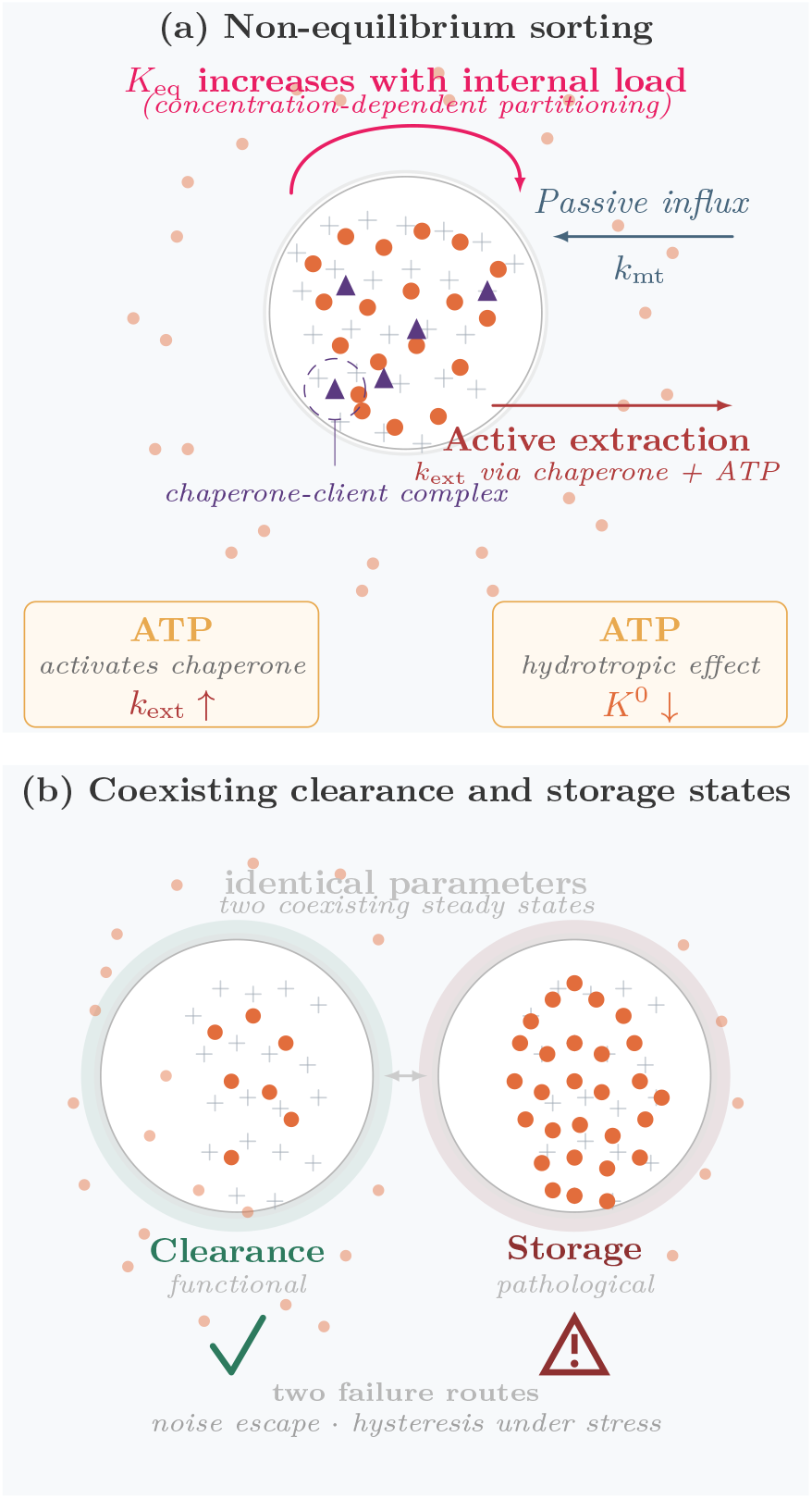
Non-equilibrium sorting and the bistable fate of condensates. (a) The condensate exchanges clients with the reservoir through two competing fluxes, a passive influx (*k*_mt_) and an ATP-driven active extraction by chaperones (*k*_ext_). The dashed circle marks a chaperone-client complex. The magenta arrow denotes concentration-dependent partitioning, *K*_eq_ increasing with internal load. The two lower panels show the dual role of ATP, chaperone activation (*k*_ext_ ↑) and hydrotropic suppression of baseline affinity (*K*^0^ ↓). The arrow marks the condition Da_0_ *< e*^*−*2^. (b) For identical parameters, the same condensate admits two coexisting steady states, functional clearance on the left and pathological storage on the right. The two states interconvert via noise-driven escape or quasi-static hysteresis under cyclic stress.

**FIG. 2.**
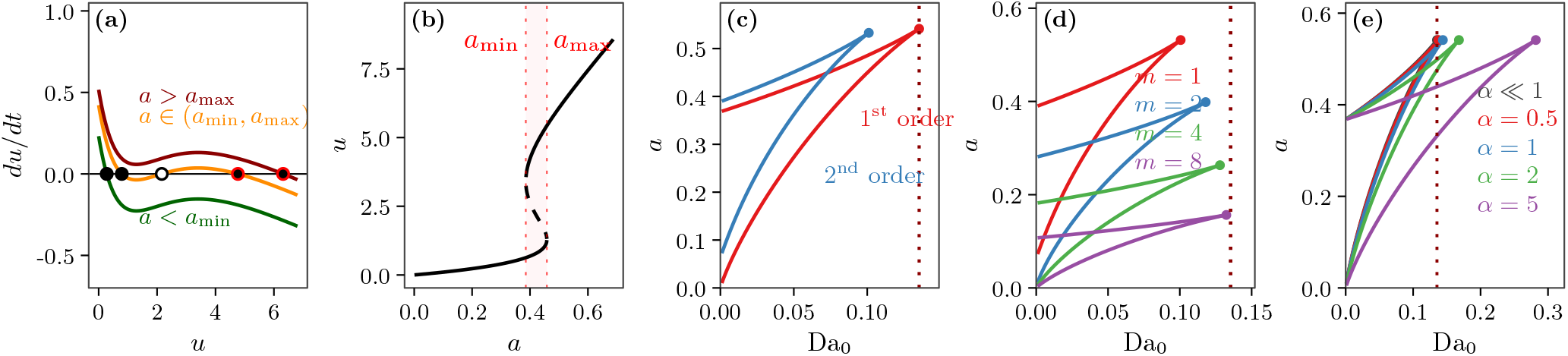
Bistability and robustness of the active sorting threshold. (a) Phase portrait *du/dt* versus *u* for three external supply levels *a*. (b) Bifurcation diagram *u* versus *a*. Solid and dashed lines are stable and unstable branches. The pink region marks the bistable window *a*_min_ *< a < a*_max_. (c) Da_0_ versus *a* for first-order and second-order feedback. The quadratic saturation term shifts the cusp leftward from *e*^*−*2^. (d) Effect of the number of client species *m*, treated within the dominant-mode approximation *λ*_max_ = *γ*_0_ + (*m* − 1)*g*. Increasing *m* strengthens the collective feedback and moves the cusp toward the ideal value *e*^*−*2^. (e) Effect of internal concentration gradients. The effectiveness factor *η*(*α*) reduces clearance and shifts the boundary to larger Da_0_ as the Thiele modulus *α* increases. The homogeneous limit *α* ≪ 1 recovers *e*^*−*2^. Parameters are *γ*_0_ = 1, *β*_0_ = 0.05, and *g* = 0.35 in (d).

Corrections from uniform internal concentration, linear extraction kinetics, and pure exponential feedback shift the cusp away from the ideal position Da_0_ = *e*^*−*2^.

Including the quadratic saturation term,

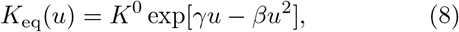

where *β >* 0 is defined microscopically in Supplemental Material Sec. S2.A, bends the feedback downward at high concentrations and lowers the critical clearance to *e*^*−*2^(1 − 4*β/γ*^2^). The phase diagram in Fig. 2(c) compares the first-order and second-order boundaries, showing the cusp move leftward. Client chain length *N* plays a similar role. In the Flory-Huggins theory both *γ* and *β* scale with *N*, which amplifies the positive feedback while strengthening saturation. Fig. 2(d) shows this systematic shift of the cusp with *N*.

Internal concentration gradients present a different correction. The Thiele modulus 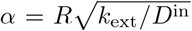 compares the droplet radius to the distance a client diffuses before extraction. When *α* is not negligible, extraction occurs predominantly near the surface and the overall clearance rate is reduced by the effectiveness factor

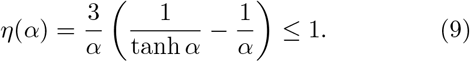

The phase diagram in Fig. 2(e) traces this effect for several values of *α*. The homogeneous limit *α* ≪ 1 recovers the ideal threshold.

These corrections can be approximately combined into a heuristic effective Damköhler number

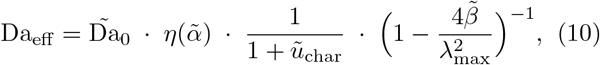

where *λ*_max_ is the largest eigenvalue of the client-client feedback matrix, 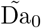 is the dominant-mode average of the zero-concentration Damköhler numbers, 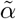 is built from the corresponding average extraction rate, 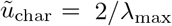, and 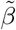 is the dominant-mode saturation coefficient, see Supplemental Material Sec. S2.C for its derivation and limitations. This expression is a heuristic guide, not a rigorous factorization for arbitrary parameters. The bifurcation condition then retains the form

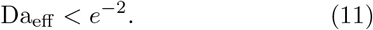

In the bare exponential-feedback model, the critical condition is exactly Da_0_ *< e*^*−*2^. Microscopic corrections shift the numerical value of the critical point by (1) amounts, but the saddle-node topology and the form of the threshold Da_eff_ *< e*^*−*2^ are preserved.

### Functional lifetime under molecular noise

The bistability of Eq. (6) is a mean-field prediction. Real condensates are finite, and the discrete nature of chemical reactions generates intrinsic noise. In the low-load limit, this noise is approximately additive and Gaussian, with intensity Λ ∝ 1*/*(*V K*_*M*_), where *V* is the condensate volume and *K*_*M*_ is the Michaelis constant, see Supplemental Material Sec. S3 for a derivation from Poisson statistics. Thus Λ scales inversely with the droplet volume as Λ ∝ 1*/R*^3^. This noise can drive stochastic transitions from the clearance state to the storage state, so the clearance state has a finite functional lifetime.

The deterministic dynamics are governed by an effective potential 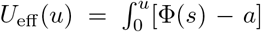 *ds*, which has a double-well shape in the bistable regime. Supplemental Material Sec. S3 shows how the stochastic dynamics map onto an overdamped Langevin equation in this potential. In the weak-noise limit, the one-dimensional projection onto the slow concentration variable *u* admits an effective Boltzmann description with temperature *τ*_0_Λ, where *τ*_0_ ≡*K*^0^*/k*_mt_ is the intrinsic relaxation time of the concentration dynamics. This approximation holds when the system remains near the deterministic fixed points. For the parameter ranges considered here, the condition *τ*_0_Λ ≪ Δ*U*_for_ is satisfied, see Supplemental Material Sec. S3.B for the Ginzburg criterion. Under this approximation, the mean first-passage time from the clearance well to the storage well follows from Kramers’ escape theory,

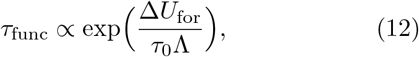

where Δ*U*_for_ = *U*_eff_ (*u*_barr_) − *U*_eff_ (*u*_low_) is the forward barrier height. The Kramers formula is valid for *τ*_0_Λ ≪ Δ*U*_for_. Near the saddle-node bifurcation, where Δ*U*_for_ → 0, the formula breaks down. The scaling laws derived below are therefore asymptotic predictions that require numerical verification in the critical region, see Fig. 5.

The barrier height depends sensitively on the clearance strength Da_0_. Near the cusp Da_0_ = *e*^*−*2^ the barrier collapses quadratically as Δ*U*_for_ ∝ (*e*^*−*2^ − Da_0_)^2^, so the functional lifetime drops exponentially. This scaling follows from expanding Φ(*u*) around the inflection point *u*_*c*_ = 2*/γ*, which yields the cubic normal form *B* = (*e*^*−*2^*/*6)*x*^3^ − *εx* with *x* = *γu* − 2 and *ε* = *e*^*−*2^ − Da_0_, see Supplemental Material Sec. S3.E. Because the effective noise strength *τ*_0_Λ ∝ 1*/R*^3^, see Supplemental Material Sec. S3.A, larger condensates are exponentially more stable against this noise-driven failure, with the stability scaling cubically in the droplet radius.

The parameter Da_0_ is itself a non-monotonic function of ATP. Two competing effects dominate. ATP activates chaperones, which increases *k*_ext_, and acts as a hydrotrope, which reduces the baseline affinity *K*^0^. The net effect is a unimodal Da_0_(*c*_*T*_), whose maximum is typically of order *e*^*−*2^ or larger. At very low ATP, the barrier is large and the clearance state is kinetically long-lived. At high ATP, if the hydrotropic suppression does not reduce Da_0_ below *e*^*−*2^, then Da_0_ *> e*^*−*2^, the storage state disappears, and clearance is permanently guaranteed. If, however, the hydrotropic branch drives Da_0_ below *e*^*−*2^ again at very high ATP, the system can re-enter the bistable regime and a second vulnerable window may appear. The model therefore predicts at least one window of maximal vulnerability when Da_0_(*c*_*T*_) crosses *e*^*−*2^ from above. Additional windows occur if the unimodal Da_0_(*c*_*T*_) crosses *e*^*−*2^ more than once. At intermediate ATP, however, Da_0_ falls just below *e*^*−*2^. The barrier collapses and even small fluctuations can trigger spontaneous contamination. The model thus predicts a window of maximal vulnerability under partial metabolic stress. For typical values *R* ~ 1 *µ*m, *D*^out^ ~ 10 *µ*m^2^*/*s, and *k*_ext_ ~ 0.1 s^*−*1^, one finds Da_0_ ∼ 10^*−*2^–10^*−*1^, which can indeed lie below *e*^*−*2^ for moderate baseline affinities, placing the system in the bistable regime. The non-monotonic ATP dependence of Da_0_ and the resulting collapse of the functional lifetime are summarized in Fig. 3(a,b). As shown in Fig. 3(c), the simulated data collapse onto the analytical prediction with slope unity, which confirms the exponential dependence of *τ*_func_ on the barrier height. The model also predicts ln *τ*_func_ ∝ *R*^3^Δ*U*_for_, see Supplemental Material Sec. S3.A, so larger condensates are exponentially more stable.

**FIG. 3.**
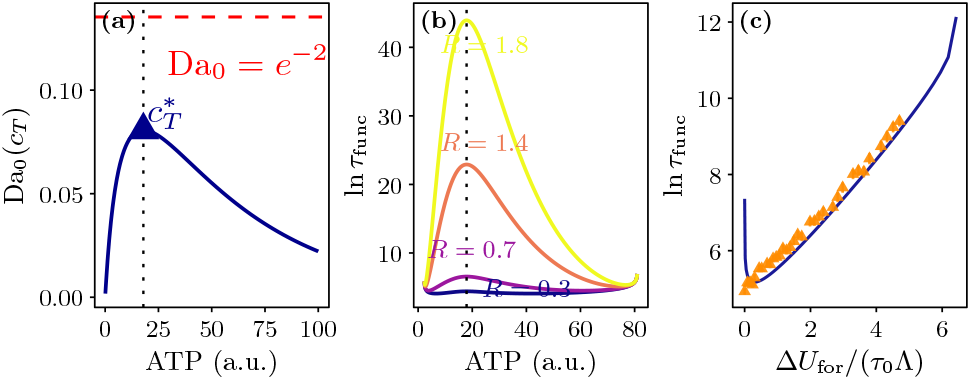
Functional lifetime and its non-monotonic control. (a) Clearance strength Da_0_ versus ATP concentration *c*_*T*_, combining Michaelis-Menten activation and hydrotropic suppression. The red dashed line marks Da_0_ = *e*^*−*2^ and the blue triangle marks the ATP concentration 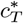 where Da_0_ is maximal. (b) ln *τ*_func_ versus ATP for several droplet radii *R*. As Da_0_ approaches *e*^*−*2^ from below, the barrier collapses and *τ*_func_ drops sharply, creating a window of maximal vulnerability at intermediate ATP. (c) Kramers scaling test. ln *τ*_func_ versus Δ*U*_for_*/*(*τ*_0_Λ). The blue line is the analytical prediction with slope unity and orange triangles are chemical Langevin simulations. Error bars are standard errors. Parameters are *γ* = 1, *τ*_0_ = 1, and Λ = 0.001.

### Energetic cost of recovery and hysteresis

*—*Once a condensate occupies the storage state, passive recovery becomes exceedingly slow. The reverse barrier Δ*U*_rev_ is typically large. A detailed calculation is given in Supplemental Material Sec. S4. Cells do not wait for a rare fluctuation. They reset the condensate actively by upregulating chaperones. This process consumes ATP. Within the effective equilibrium description defined by the potential *U*_eff_ and temperature *τ*_0_Λ, the Crooks fluctuation theorem applies to protocols that change Da_0_ quasi-statically, which gives the bound

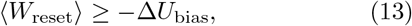

where Δ*U*_bias_ = *U*_eff_ (*u*_high_) − *U*_eff_ (*u*_low_). This bound relies on the effective-equilibrium assumption, which is approximate for an active system. Without this assumption, a weaker but more general bound follows from the second law of thermodynamics, ⟨*W*_reset_⟩ ≥ −Δ*F*, with Δ*F* the free-energy difference between the two effective states, see Supplemental Material Sec. S4.B. The two bounds coincide only when the effective-equilibrium description is valid. For weak basal chaperone activity, Da_0_ ≪ *e*^*−*2^, the free-energy bias depends on the path taken in the (Da_0_, *a*) plane. If the external supply *a* is held fixed, the storage state moves to ever larger concentrations as *u*_high_ ≃ *a/*Da_0_, and the bias diverges as Δ*U*_bias_ ≃ − *a*^2^*/*(2Da_0_). Thus the resetting cost does not saturate; it grows without bound as chaperone activity vanishes. Condensates that are very stable against noise become effectively irreversible and increasingly expensive to rescue.

Cells also face cyclic challenges. During a stress episode the external supply *a* of misfolded clients rises slowly and then returns to basal levels. If this variation is slow compared to internal relaxation, the system tracks the steady-state branches of the bifurcation diagram. It executes an irreversible hysteresis loop. The energy dissipated per full cycle must be supplied by ATP hydrolysis. Near the cusp it scales quadratically with the distance to the critical point,

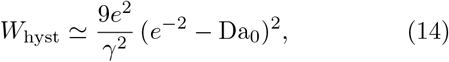

as shown in Supplemental Material Sec. S4.C. This quadratic scaling means the hysteresis loss vanishes smoothly at the cusp, which reflects the collapse of the bistable window. The corresponding forward-barrier scaling is Δ*U*_for_ ∝ (*e*^*−*2^ − Da_0_)^2^, derived in Supplemental Material Sec. S3.E and displayed in Fig. 4(a). Both the resetting work and the hysteresis loss, when expressed in physical units, are proportional to the condensate volume *V* ∝ *R*^3^. The dimensionless energies above are converted to physical free energies by the factor *k*_B_*T* (*V K*_*M*_)*/a*, where *V* ∝ *R*^3^ is the droplet volume, *K*_*M*_ is the Michaelis constant, and *a* is the dimensionless external supply. Hence the physical resetting and hysteresis costs scale as *R*^3^. This gives a physical reason for the upper size limit of functional organelles. Droplets that grow too large become metabolically unaffordable to maintain and rescue. The quadratic collapse of the forward barrier and the hysteresis area are shown in Fig. 4(a,b). Panel (c) shows the free-energy bias for fixed external supply *a* = 0.1. As Da_0_ → 0, the bias diverges as |Δ*U*_bias_| ≃ *a*^2^*/*(2Da_0_), confirming that weak chaperone activity makes the storage state energetically inaccessible and the resetting cost prohibitively large.

**FIG. 4.**
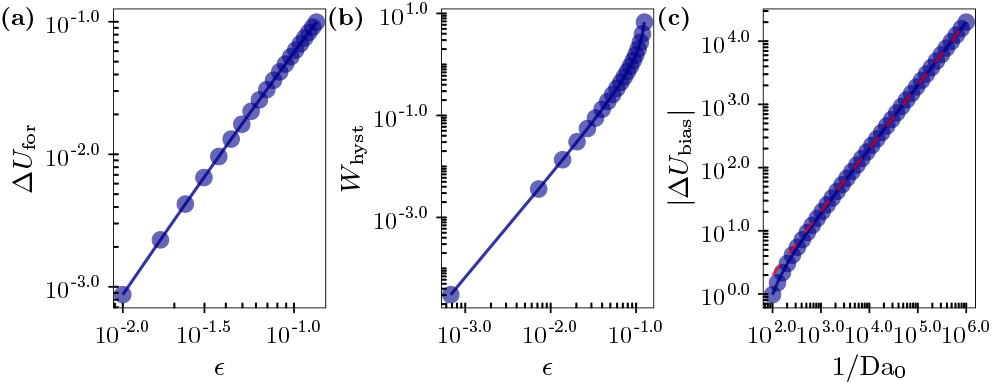
Energetic cost of recovery and hysteresis. (a) Log-log plot of Δ*U*_for_ versus *ε* = *e*^*−*2^ − Da_0_. The blue line shows Δ*U*_for_ ∝ *ε*^2^. (b) Log-log plot of the hysteresis area *W*_hyst_ versus *ε*. The numerical integration follows *W*_hyst_ ≃ (9*e*^2^*/γ*^2^)*ε*^2^. (c) Free-energy bias |Δ*U*_bias_| versus Da_0_. The blue curve shows Δ*U*_bias_| → 2*e*^*−*2^*/γ*^2^ as Da_0_ → 0, with a subleading correction Da_0_ ln(Da_0_*/e*^*−*2^). All quantities are dimensionless. The energy scale is *k*_*B*_*T* (*V K*_*M*_)*/a*, where *V* is the condensate volume and *a* is the dimensionless external supply. Parameters are *γ* = 1, *τ*_0_ = 1, and Λ = 0.001.

### Critical fluctuations and early warning signals

*—*The saddle-node bifurcation is a critical point. Its approach is marked by characteristic dynamical signatures that can be detected while the condensate is still in the functional, monostable regime. Linearizing the stochastic dynamics around the clearance fixed point yields an Ornstein-Uhlenbeck process. Its steady-state variance and relaxation time are

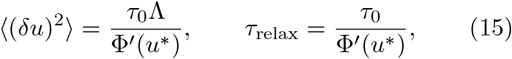

where Φ′(*u*^*\**^) is the slope of Φ(*u*) at the fixed point, *τ*_0_ is the intrinsic relaxation time of the concentration dynamics, and *τ*_0_Λ plays the role of an effective temperature in the weak-noise limit, see Supplemental Material Sec. S5. As the bifurcation is approached, Φ′(*u*^*\**^) → 0, and both the variance and the relaxation time diverge.

The scaling exponents depend on the path taken in the (Da_0_, *a*) phase diagram, see Fig. 5(a). These are meanfield exponents of the saddle-node bifurcation. The system does not exhibit a genuine critical point with diverging correlation length.

**FIG. 5.**
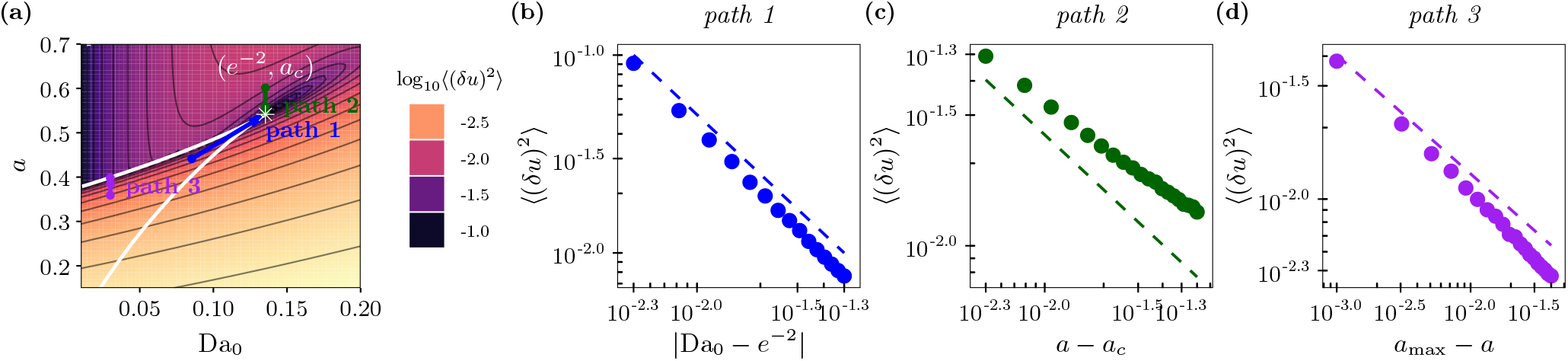
Critical fluctuations and scaling near the saddle-node bifurcation. (a) Steady-state variance log_10_⟨(*δu*)^2^⟩ in the (Da_0_, *a*) plane from the linear-noise approximation. The white star marks the cusp (Da_0*c*_, *a*_*c*_) = (*e*^*−*2^, 4*/γe*^2^). Three paths are tested and labeled in panel (a), path 1 along *a* = *a*_*c*_ − 2(*e*^*−*2^ − Da_0_)*/γ* in blue, path 2 along *a* at fixed Da_0_ = *e*^*−*2^ in green, and path 3 approaching the upper saddle-node boundary at Da_0_ = 0.03 in purple. (b)-(d) Log-log plots of variance versus distance to the bifurcation for the three paths. Points are Monte Carlo samples of the linearized Ornstein-Uhlenbeck dynamics with 100 independent trajectories per point. Error bars are standard errors. Colored dashed lines show the asymptotic power laws with exponents −1, −2*/*3, and −1*/*2, and the corresponding slopes are annotated inside each panel. Parameters are *γ* = 1, Λ = 0.001, and *τ*_0_ = 1.

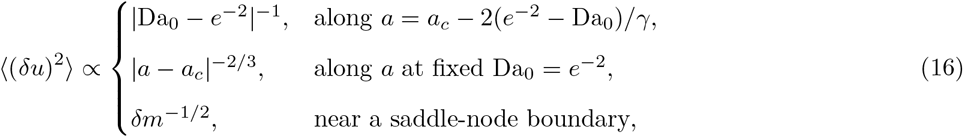

where in the first case the supply is tuned to keep the fixed point at the inflection point *u*^*\**^ = 2*/γ*. In the last line *δm* denotes the distance to the boundary in the parameter being varied, for example *δm* = *a*_max_ − *a* or *δm* = |Da_0_ − Da_0,max_|. The relaxation time *τ*_relax_ obeys identical scaling laws.

These linear scalings hold when the noise is sufficiently weak that the cubic nonlinearity of the potential remains negligible. The quantitative condition is given in Supplemental Material Sec. S5.B. The three scaling paths and the corresponding exponents are tested in Fig. 5(b)– (d). As shown there, the simulated steady-state variances follow the theoretical power laws with exponents −1, −2*/*3, and −1*/*2, respectively, which confirms the path-dependent critical divergence.

These signatures are directly measurable. Fluorescence correlation spectroscopy (FCS) records the steady-state fluctuation amplitude non-invasively. Fluorescence recovery after photobleaching (FRAP) probes the relaxation time. In a reconstituted system one may titrate a chaperone inhibitor to reduce Da_0_ while monitoring the FCS amplitude and FRAP half-time. A gradual rise in the FCS amplitude and a simultaneous lengthening of the FRAP recovery half-time provide an early warning signal that appears well before the condensate transitions to the pathological storage state.

## Discussion

*—*We have shown that active client extraction can be reduced to a single algebraic equation. The ideal exponential-feedback limit yields a threshold Da_0_ = *e*^*−*2^ that does not depend on the feedback strength. Corrections from internal gradients, Michaelis-Menten saturation, and higher-order thermodynamic terms shift this threshold but preserve the saddle-node bifurcation topology. The model gives a closed-form criterion linking molecular parameters such as the baseline partition coefficient *K*^0^, the chaperone extraction rate *k*_ext_, and the diffusive supply *k*_mt_ to condensate stability.

The MNSR model offers an analytically tractable limiting case of field-theoretic descriptions for chemically active emulsions and active Cahn-Hilliard systems [19, 21– 25, 33, 34]. The sharp threshold *e*^*−*2^ is not readily visible in numerical phase diagrams, yet it emerges from the algebraic structure and can serve as a benchmark for more detailed simulations. The model also makes several experimentally testable predictions. Selectivity arises through Da_0_, which combines *K*^0^, *k*_ext_, and *k*_mt_. Two clients with identical *K*^0^ but different *k*_ext_ can reside on opposite sides of the *e*^*−*2^ threshold, which offers a quantitative rationale for selective extraction observed in nucleoli and stress granules [8, 9, 11]. Disease-linked mutations that strengthen homotypic interactions increase *γ*, narrow the safe operating regime, and are consistent with accelerated liquid-to-solid transitions in FUS and TDP-43 [4–7, 35–37]. The predicted non-monotonic ATP dependence implies a window of maximal vulnerability at intermediate ATP, which may explain why partial energy depletion is often more damaging than complete ATP loss in neurodegenerative contexts [29–32, 38].

Several limitations should be noted. We assumed a spatially uniform intra-condensate concentration and a passive, well-mixed chaperone pool. Real condensates may exhibit internal gradients, cooperative chaperone binding, and interface-localized extraction [12, 16, 18, 39]. The robustness of the *e*^*−*2^ threshold to the corrections we considered suggests that the bifurcation scenario persists under more realistic descriptions. Experimentally, the key parameters *K*^0^, *k*_ext_, and *D*^out^ can be measured by established fluorescence methods [39– 42]. FCS and FRAP can monitor the predicted critical slowing down near the saddle-node boundary, and the quadratic scaling of hysteresis loss and the *R*^3^ scaling of recovery cost can be tested by varying droplet size. These signatures provide early warning signals for condensate dysfunction and establish the MNSR model as a quantitative framework for condensate quality control. Engineered chaperones that modulate condensate stability [43, 44] offer a promising route to test these predictions therapeutically.

G.Y. acknowledges support from the National Natural Science Foundation of China (Grant No. 32001242).

