## Supplementary material for "*e*^−2^ Threshold for Bistability in Non-equilibrium Sorting of Biomolecular Condensates": suppl

This Supplemental Material provides the detailed derivation of the Minimal Non-equilibrium Sorting Reactor (MNSR) model, the thermodynamic origin of concentration-dependent partitioning, the saddle-node bifurcation analysis, the derivation of the effective Langevin dynamics, the functional lifetime, the energetic cost of recovery, and the critical fluctuations. Equation and figure numbers are prefixed with "S" to distinguish them from the main text.

### DERIVATION OF THE MNSR MODEL

In this section we derive the Minimal Non-equilibrium Sorting Reactor (MNSR) equations from a multi-component Flory-Huggins free energy augmented by active extraction and diffusive transport. We begin by specifying the free energy functional and the exchange chemical potentials. We then formulate the reaction-diffusion dynamics, take the sharp-interface limit with the Laplace pressure correction, perform an internal homogenisation when diffusion is fast, apply the quasi-steady approximation for external diffusion to obtain a mass-transfer coefficient, and finally write the steady-state mass balance in dimensionless form.

#### Free energy of a multi-component mixture

We consider a system composed of a scaffold protein $S$, $n$ client species $C_i$ ($i = 1, \ldots, n$), and solvent $W$. The mixture is taken as incompressible, so the volume fractions obey

$$\phi_S(\mathbf{r}, t) + \sum_{i=1}^{n} \phi_i(\mathbf{r}, t) + \phi_W(\mathbf{r}, t) = 1 \tag{S1}$$

at every point in space. The total free energy (in units of thermal energy $k_BT$) is a functional of the volume-fraction fields,

$$\frac{F[\phi_S, \{\phi_i\}]}{k_BT} = \int_V \left[ f_{\text{mix}}(\{\phi\}) + \frac{\ell_S}{2}|\nabla\phi_S|^2 + \sum_{i=1}^{n} \frac{\ell_i}{2}|\nabla\phi_i|^2 \right] dV. \tag{S2}$$

The first term $f_{\text{mix}}$ is the Flory-Huggins mixing free-energy density,

$$\begin{aligned} f_{\text{mix}} = {} & \frac{\phi_S}{N_S}\ln\phi_S + \sum_{i=1}^{n} \frac{\phi_i}{N_i}\ln\phi_i + \phi_W \ln\phi_W \\ & + \chi_{SW}\phi_S\phi_W + \sum_{i=1}^{n} \chi_{Si}\phi_S\phi_i + \sum_{i=1}^{n} \chi_{iW}\phi_i\phi_W + \sum_{i<j} \chi_{ij}\phi_i\phi_j. \end{aligned} \tag{S3}$$

Here $N_S$ and $N_i$ are the chain lengths (numbers of lattice sites) relative to a solvent molecule, and the $\chi_{\alpha\beta}$ are dimensionless interaction parameters. The gradient terms, with coefficients $\ell_S, \ell_i > 0$, represent the free-energy cost of concentration inhomogeneities and determine the interfacial tensions.

Because of incompressibility, the independent fields are $\phi_S$ and $\{\phi_i\}$ while $\phi_W$ is slaved by Eq. (S1). The chemical potential of species $k$ (in units of $k_BT$) is obtained by functional differentiation,

$$\mu_k(\mathbf{r}) = \frac{\delta(F/k_BT)}{\delta\phi_k(\mathbf{r})}, \tag{S4}$$

and the exchange chemical potential with respect to the solvent is defined as

$$\tilde{\mu}_k = \mu_k - \mu_W. \tag{S5}$$

For a client protein $i$, neglecting gradient terms for the moment, the exchange chemical potential can be written as

$$\tilde{\mu}_i = \frac{1}{N_i}\ln\frac{\phi_i}{\phi_W} - \frac{1}{N_S}\ln\phi_S - (\chi_{Si}-\chi_{iW})\phi_S + \sum_j \chi_{ij}\phi_j + \text{const.} \tag{S6}$$

In Eq. (S6) we have omitted terms that are linear in the volume fractions and can be absorbed into the constant. The precise form of the constant is not needed for the subsequent analysis.

For the purpose of calculating the equilibrium partition coefficient of a client between the dense and dilute phases of the scaffold, it is sufficient to use the simplified form

$$\tilde{\mu}_i \simeq \frac{1}{N_i}\ln\phi_i - \ln\phi_W - (\chi_{Si}-\chi_{iW})\phi_S + \sum_j \chi_{ij}\phi_j + \text{const}, \tag{S7}$$

which captures the leading dependence on $\phi_i$, $\phi_W$, and the scaffold-client interaction $\chi_{Si}$. Equation (S7) is obtained from Eq. (S6) by dropping terms that are constant or proportional to $\phi_S$ in the dilute external phase (where $\phi_S^{\text{out}} = 0$) and by absorbing the solvent interaction into an effective constant.

**Reaction-diffusion dynamics with active extraction**

The time evolution of the conserved volume fractions follows the Cahn-Hilliard-Cook equations with an additional reaction term that breaks detailed balance,

$$\frac{\partial\phi_S}{\partial t} = \nabla\cdot\big(M_S\nabla\tilde{\mu}_S\big), \tag{S8}$$

$$\frac{\partial\phi_i}{\partial t} = \nabla\cdot\big(M_i\nabla\tilde{\mu}_i\big) + \mathcal{R}_i(\{c_j\}, c_T). \tag{S9}$$

The scaffold mobility $M_S$ and client mobilities $M_i$ are related to the respective diffusion coefficients by $D_S = M_S/N_S$ and $D_i = M_i/N_i$. The term $\mathcal{R}_i$ accounts for the ATP-driven extraction of client $i$ by molecular chaperones. Because chaperones are large and slowly diffusing, we assume that extraction takes place only inside the scaffold-rich phase.

To connect the reaction kinetics, conveniently expressed in terms of molar concentrations, with the evolution of volume fractions, we introduce the partial molecular volumes $v_i$ and write $\phi_i = v_i c_i$, where $c_i$ is the molar concentration of client $i$ (an analogous relation holds for the scaffold, but its explicit concentration is not required in the extraction term). A simple model with competitive Michaelis-Menten kinetics is then

$$\mathcal{R}_i(\mathbf{r},t) = -\,\Theta(\phi_S-\phi_{\text{th}})\, v_i k_{\text{cat},i} E_{\text{act}}^{\text{in}} \frac{c_i/K_{M,i}}{1+\sum_j c_j/K_{M,j}}. \tag{S10}$$

Here $\Theta(\cdots)$ is the Heaviside step function (with a small threshold $\phi_{\text{th}}$), $k_{\text{cat},i}$ is the catalytic rate constant, $K_{M,i}$ is the Michaelis constant for client $i$ (with dimensions of concentration), and $E_{\text{act}}^{\text{in}}$ is the concentration of active chaperone inside the dense phase. The denominator $1+\sum_j c_j/K_{M,j}$ reflects the competition of different clients for the same chaperone pool. The factor $v_i$ converts the molar reaction rate into the rate of change of volume fraction, ensuring dimensional consistency.

The active chaperone concentration depends on the availability of ATP. Assuming rapid equilibrium for ATP binding to the chaperone,

$$E_{\text{act}}^{\text{in}} = E_{\text{tot}}^{\text{in}} \frac{c_T}{K_{M,\text{ATP}} + c_T}, \tag{S11}$$

where $c_T$ is the ATP concentration and $K_{M,\text{ATP}}$ is the ATP Michaelis constant. The total chaperone concentration inside the condensate, $E_{\text{tot}}^{\text{in}}$, is itself determined by the chaperone's own equilibrium partition coefficient.

When the client load is low, $\sum_j c_j/K_{M,j} \ll 1$, the extraction kinetics simplifies to a first-order reaction in the client concentration. Using $\phi_i = v_i c_i$, we obtain

$$\mathcal{R}_i \approx -\,\Theta(\phi_S-\phi_{\text{th}})\, k_{\text{ext},i}\phi_i, \qquad k_{\text{ext},i} \equiv \frac{k_{\text{cat},i}E_{\text{act}}^{\text{in}}}{K_{M,i}}. \tag{S12}$$

The extraction rate constant $k_{\text{ext},i}$ (units of inverse time) condenses the entire chaperone activity into a single parameter. It depends on ATP both through the Michaelis-Menten activation in Eq. (S11) and, indirectly, through possible ATP-dependent changes in $E_{\text{tot}}^{\text{in}}$.

#### Sharp-interface limit and Laplace pressure correction

In the absence of clients, the scaffold-solvent binary mixture described by the free energy $f_0(\phi_S) = f_{\text{mix}}(\phi_S, \phi_i{=}0)$ can phase separate when $\chi_{SW}$ exceeds a critical value. The equilibrium volume fractions of the coexisting phases, $\phi_S^{\text{in}}$ (dense) and $\phi_S^{\text{out}}$ (dilute), are found from the common-tangent construction on $f_0(\phi_S)$. For a spherical droplet of radius $R$ much larger than the interfacial width $w_{\text{int}} \sim \sqrt{\ell_S/f_0''}$, the scaffold profile can be approximated by a sharp interface,

$$\phi_S(\mathbf{r}) = \begin{cases} \phi_S^{\text{in}}, & |\mathbf{r}| < R, \\ 0, & |\mathbf{r}| > R, \end{cases} \tag{S13}$$

where we have used $\phi_S^{\text{out}} \approx 0$ for a sufficiently asymmetric phase diagram.

The presence of a curved interface generates a Laplace pressure difference

$$\Delta p = p^{\text{in}} - p^{\text{out}} = \frac{2\sigma}{R}, \tag{S14}$$

with the interfacial tension $\sigma$ given by the classical formula

$$\sigma = \int_{\phi_S^{\text{out}}}^{\phi_S^{\text{in}}} \sqrt{2\ell_S\big[f_0(\phi) - f_0(\phi_S^{\text{out}}) - \mu_S^{\text{eq}}(\phi - \phi_S^{\text{out}})\big]}\, d\phi. \tag{S15}$$

The equilibrium condition for a client protein across the interface is the continuity of the full chemical potential, which includes the pressure-volume work,

$$\tilde{\mu}_i^{\text{in}} + \frac{v_i p^{\text{in}}}{k_B T} = \tilde{\mu}_i^{\text{out}} + \frac{v_i p^{\text{out}}}{k_B T}, \tag{S16}$$

where $v_i$ is the partial molecular volume of client $i$.

To obtain the partition coefficient, we insert Eq. (S7) into Eq. (S16). In the external dilute phase we have $\phi_S^{\text{out}} = 0$, $\phi_i^{\text{out}} \ll 1$, $\phi_W^{\text{out}} \approx 1$, and $\sum \chi_{ij}\phi_j^{\text{out}} \approx 0$. In the internal dense phase we write $\phi_W^{\text{in}} = 1 - \phi_S^{\text{in}} - \sum_j \phi_j^{\text{in}} \approx 1 - \phi_S^{\text{in}}$, where we neglect the small client volume fractions inside the condensate for the baseline partitioning. Collecting terms gives

$$\frac{1}{N_i} \ln \frac{\phi_i^{\text{in}}}{\phi_i^{\text{out}}} = \ln(1 - \phi_S^{\text{in}}) + (\chi_{Si} - \chi_{iW})\phi_S^{\text{in}} - \frac{v_i}{k_B T}\frac{2\sigma}{R} + \mathcal{O}(\phi_i^{\text{in}}). \tag{S17}$$

Exponentiating and defining the partition coefficient $K_{\text{eq},i} \equiv \phi_i^{\text{in}}/\phi_i^{\text{out}}$, we obtain

$$K_{\text{eq},i}(R, c_T) = K_{\text{eq},i}^0(c_T)\, \exp\Big( - N_i \frac{v_i}{k_B T}\frac{2\sigma}{R}\Big), \tag{S18}$$

Here $v_i$ denotes the monomeric volume of client $i$, so that the full-chain volume is $N_i v_i$. Equivalently, if $v_i$ is instead defined as the full-chain molecular volume, then the pressure term in Eq. (S16) and the exponent in Eq. (S17) must be divided by $N_i$, and the Laplace factor in Eq. (S18) remains $\exp[-v_i(2\sigma)/(k_B T R)]$. The baseline equilibrium coefficient is

$$K_{\text{eq},i}^0(c_T) = \big(1 - \phi_S^{\text{in}}\big)^{N_i}\, \exp\Big[N_i\big(\chi_{Si}(c_T) - \chi_{iW}\big)\phi_S^{\text{in}}\Big]. \tag{S19}$$

Equation (S19) shows how the scaffold-client affinity (encoded in $\chi_{Si}$) and the ATP concentration (which can modulate $\chi_{Si}$ through hydrotropic effects) determine the baseline enrichment. The exponential factor in Eq. (S18) is the Laplace correction, which becomes important for radii $R \lesssim 2\sigma v_i/(k_B T)$. In the remainder of this paper we absorb the Laplace term into a renormalised $K_{\text{eq},i}$ and treat the partition coefficient as a prescribed function of $c_T$ and, later, of the client concentrations themselves (owing to $\chi_{ij} \neq 0$).

#### Internal homogenisation and the Thiele modulus

Inside the droplet, the client concentration $c_i^{\text{in}}$ (related to $\phi_i^{\text{in}}$ by $\phi_i^{\text{in}} = v_i c_i^{\text{in}}$) obeys a reaction-diffusion equation. Substituting the sharp interface profile (S13) into Eq. (S9) and using the dilute-client extraction rate (S12), we obtain, for $|\mathbf{r}| < R$,

$$\frac{\partial c_i^{\text{in}}}{\partial t} = D_i^{\text{in}} \nabla^2 c_i^{\text{in}} - k_{\text{ext},i} c_i^{\text{in}}, \tag{S20}$$

where $D_i^{\rm in} = M_i/N_i$ is the translational diffusion coefficient inside the dense phase.

Equation (S20) introduces a characteristic length scale $L_i \equiv \sqrt{D_i^{\rm in}/k_{{\rm ext},i}}$, called the clearance length, which is the typical distance a client molecule diffuses before being extracted. The ratio of the droplet radius to this length defines the Thiele modulus

$$\alpha_i \equiv \frac{R}{L_i} = R\sqrt{\frac{k_{{\rm ext},i}}{D_i^{\rm in}}}. \tag{S21}$$

When $\alpha_i \ll 1$, diffusion is much faster than extraction, and the concentration field $c_i^{\rm in}$ remains essentially spatially uniform. In this reaction-limited regime we can replace $c_i^{\rm in}(\mathbf{r},t)$ by a single, time-dependent value $c_i^{\rm in}(t)$. The total extraction rate is then simply

$$\mathcal{R}_i^{\rm cons}(t) = \int_V k_{{\rm ext},i} c_i^{\rm in}\, dV = V k_{{\rm ext},i} c_i^{\rm in}(t), \qquad V = \frac{4}{3}\pi R^3. \tag{S22}$$

If $\alpha_i$ is not small, concentration gradients develop inside the droplet, and the extraction rate is reduced by an effectiveness factor $\eta(\alpha_i) < 1$. For the purpose of this work we restrict ourselves to the analytically transparent case $\alpha_i \ll 1$ because all the essential non-equilibrium physics is already present in this limit.

#### Quasi-steady external diffusion

Outside the droplet, the scaffold is absent and the extraction machinery is negligible. The client concentration $c_i^{\rm out}$ evolves by free diffusion,

$$\frac{\partial c_i^{\rm out}}{\partial t} = D_i^{\rm out}\nabla^2 c_i^{\rm out}, \tag{S23}$$

with a diffusion coefficient $D_i^{\rm out}$ that may differ from $D_i^{\rm in}$ because the dense phase is more crowded. The exterior is assumed to be a large reservoir that maintains a fixed bulk concentration $c_i^{\rm out}(\infty) = c_i^{\rm out}$ far from the droplet.

The time scale for the external diffusion field to relax is $\tau_{\rm ext} \sim R^2/D_i^{\rm out}$. If $\tau_{\rm ext}$ is short compared to the time over which the droplet composition changes, we can invoke the quasi-steady approximation, in which the external concentration profile instantly adapts to the slowly varying surface concentration. Setting $\nabla^2 c_i^{\rm out} \approx 0$ and solving the resulting Laplace equation with spherical symmetry yields the concentration profile that satisfies the boundary conditions $c_i^{\rm out}(R) = c_i^{\rm in}/K_{{\rm eq},i}$ and $c_i^{\rm out}(\infty) = c_i^{\rm out}$. This profile is

$$c_i^{\rm out}(r) = c_i^{\rm out} - \frac{R}{r}\Big(c_i^{\rm out} - \frac{c_i^{\rm in}}{K_{{\rm eq},i}}\Big), \qquad r \geq R. \tag{S24}$$

The diffusive flux at the droplet surface is obtained from Fick's law,

$$J_i^{\rm in} = -4\pi R^2 D_i^{\rm out}\left.\frac{dc_i^{\rm out}}{dr}\right|_{r=R} = 4\pi R D_i^{\rm out}\Big(c_i^{\rm out} - \frac{c_i^{\rm in}}{K_{{\rm eq},i}}\Big). \tag{S25}$$

To express this flux in a form that is symmetric with the volumetric consumption rate inside the droplet, we introduce the external mass-transfer coefficient

$$k_{{\rm mt},i} \equiv \frac{3D_i^{\rm out}}{R^2}, \tag{S26}$$

so that the influx becomes

$$J_i^{\rm in} = V k_{{\rm mt},i}\Big(c_i^{\rm out} - \frac{c_i^{\rm in}}{K_{{\rm eq},i}}\Big). \tag{S27}$$

The inverse $k_{{\rm mt},i}^{-1}$ is the characteristic time for a client molecule outside the droplet to be replenished by diffusion, and $1/k_{{\rm mt},i}$ plays the role of an external mixing time.

#### MNSR equations

At a non-equilibrium steady state, the diffusive influx must exactly balance the reactive consumption for each client, leading to

$$J_i^{\rm in} = \mathcal{R}_i^{\rm cons} \quad \Longrightarrow \quad k_{{\rm mt},i}\Big(c_i^{\rm out} - \frac{c_i^{\rm in}}{K_{{\rm eq},i}}\Big) = k_{{\rm ext},i} c_i^{\rm in}. \tag{S28}$$

It is convenient to work with dimensionless concentrations normalised by the respective Michaelis constants,

$$u_i = \frac{c_i^{\rm in}}{K_{M,i}}, \qquad u_i^{\rm out} = \frac{c_i^{\rm out}}{K_{M,i}}, \tag{S29}$$

and to introduce the Damköhler number for client $i$,

$$\mathrm{Da}_i \equiv \frac{K_{{\rm eq},i} k_{{\rm ext},i}}{k_{{\rm mt},i}}. \tag{S30}$$

$\mathrm{Da}_i$ compares the rate of active extraction (amplified by the equilibrium partition coefficient) to the rate of diffusive supply. Substituting these definitions into Eq. (S28) yields the dimensionless MNSR equation

$$K_{{\rm eq},i} u_i^{\rm out} - u_i = \mathrm{Da}_i \, \frac{u_i}{1 + \sum_j u_j}. \tag{S31}$$

The sum in the denominator accounts for saturation of the chaperone machinery when multiple clients are present at high concentrations. Equation (S31) is the central result of our coarse-graining procedure, a closed algebraic relation that determines the steady-state composition $\{u_i\}$ of the condensate in terms of the external concentrations $\{u_i^{\rm out}\}$, the partition coefficients $\{K_{{\rm eq},i}\}$, and the Damköhler numbers $\{\mathrm{Da}_i\}$.

It is useful to express $\mathrm{Da}_i$ through the two dimensionless numbers introduced above. Using $k_{{\rm mt},i} = 3D_i^{\rm out}/R^2$ and defining the diffusivity ratio $\rho_i \equiv D_i^{\rm in}/D_i^{\rm out}$, we obtain

$$\mathrm{Da}_i = K_{{\rm eq},i} \frac{\alpha_i^2 \rho_i}{3}. \tag{S32}$$

The product $\alpha_i^2 \rho_i/3$ captures the purely kinetic contribution, while $K_{{\rm eq},i}$ carries the thermodynamic information. The diffusivity ratio $\rho_i$ controls the relative importance of external versus internal transport. When $\rho_i \ll 1$, external diffusion is rapid and the system tends to be reaction-limited (small Damköhler number). When $\rho_i \gg 1$, external supply is slow and the system may enter an external-mass-transfer-limited regime. Importantly, the condition for internal homogeneity $\alpha_i \ll 1$ is not automatically guaranteed by a moderate $\mathrm{Da}_i$ and $\rho_i \lesssim 1$; it also depends on the partition coefficient through $\alpha_i = \sqrt{3\mathrm{Da}_i/(K_{{\rm eq},i}\,\rho_i)}$. For a given $\mathrm{Da}_i$ and $\rho_i$, a sufficiently large $K_{{\rm eq},i}$ ensures that $\alpha_i$ remains small. In the main text we assume $\alpha_i \ll 1$ so that the uniform-concentration MNSR equations are exact. The generalisation to non-uniform profiles via an effectiveness factor $\eta(\alpha_i)$ is discussed later.

### BISTABILITY DRIVEN BY CONCENTRATION-DEPENDENT PARTITIONING

The linear analysis assumed that the partition coefficients $K_{{\rm eq},i}$ are constants, independent of the client concentrations inside the condensate. This assumption is legitimate only when client–client interactions are negligible. In many biological condensates, however, client proteins engage in weak, multivalent homotypic and heterotypic contacts that make the free energy of the dense phase a non-linear function of the client concentrations. Such thermodynamic feedback can qualitatively alter the steady-state landscape of the MNSR equations and, as we show below, give rise to bistability and discontinuous switching between distinct functional states.

#### Thermodynamic origin of concentration-dependent partitioning

We consider a single client species X that interacts with itself, embedded in a scaffold matrix with fixed volume fraction $\phi_S^{\rm in}$ inside the dense phase. The scaffold is treated as a passive background that provides the phase-separated compartment and its concentration is taken to be constant. The relevant thermodynamic quantity is the exchange chemical potential of the client with respect to the solvent, $\tilde{\mu}_X = \mu_X - \mu_W$.

Starting from the full Flory–Huggins free energy, Eq. (S3), and the definition of exchange chemical potentials, Eq. (S5), functional differentiation gives, for a client with chain length $N$, self-interaction parameter $\chi_{XX}$, and scaffold-interaction parameter $\chi_{SX}$,

$$\begin{aligned}\tilde{\mu}_X = &\frac{1}{N}\ln\phi_X - \ln\phi_W - \frac{1}{N_S}\ln\phi_S \\ &- (\chi_{SX} - \chi_{XW})\phi_S + \chi_{XX}\phi_X - \chi_{XW}\phi_W + \text{const.}\end{aligned} \tag{S33}$$

This expression is consistent with the mean-field Flory–Huggins framework. The term proportional to $\chi_{XX}$ contains the self-interaction, and the sign convention is such that $\chi_{XX} < 0$ corresponds to an attractive effective interaction.

Inside the dense phase the scaffold volume fraction is fixed at $\phi_S = \phi_S^{\text{in}}$, and the solvent volume fraction is $\phi_W^{\text{in}} = 1 - \phi_S^{\text{in}} - \phi_X^{\text{in}}$. In the external dilute phase we have $\phi_S^{\text{out}} \approx 0$, $\phi_W^{\text{out}} \approx 1$, and $\phi_X^{\text{out}} \ll 1$. The scaffold terms and the constants are the same on both sides of the interface and cancel out when we equate chemical potentials to obtain the partition coefficient. The relevant client-dependent part of $\tilde{\mu}_X$ can therefore be written as

$$\tilde{\mu}_X(\phi_X) \simeq \frac{1}{N}\ln\phi_X - \ln(1 - \phi_S^{\text{in}} - \phi_X) + (\chi_{XX} - \chi_{XW})\phi_X + \text{const,} \tag{S34}$$

where the constant absorbs all terms that depend only on $\phi_S^{\text{in}}$ and on the external-phase composition.

For client volume fractions small compared to the available pore space, $\phi_X \ll 1 - \phi_S^{\text{in}}$, we expand the logarithmic term

$$-\ln(1 - \phi_S^{\text{in}} - \phi_X) = -\ln(1 - \phi_S^{\text{in}}) + \frac{\phi_X}{1 - \phi_S^{\text{in}}} + \frac{\phi_X^2}{2(1 - \phi_S^{\text{in}})^2} + \mathcal{O}(\phi_X^3). \tag{S35}$$

To linear order in $\phi_X$, the $\phi_X$-dependent part of $\tilde{\mu}_X$ is

$$\tilde{\mu}_X(\phi_X) \approx \frac{1}{N}\ln\phi_X + A\phi_X, \quad \text{where} \quad A \equiv \frac{1}{1 - \phi_S^{\text{in}}} + (\chi_{XX} - \chi_{XW}). \tag{S36}$$

At equilibrium across the interface the chemical potentials in the two phases are equal, and we neglect the Laplace pressure correction here as it can be absorbed into a redefinition of $K^0$. Equating $\tilde{\mu}_X(\phi_X^{\text{in}})$ and $\tilde{\mu}_X(\phi_X^{\text{out}})$ and using $\phi_X^{\text{out}} \ll 1$ yields

$$\ln\frac{\phi_X^{\text{in}}}{\phi_X^{\text{out}}} = \ln K^0 - NA\,\phi_X^{\text{in}} + \mathcal{O}\big((\phi_X^{\text{in}})^2\big), \tag{S37}$$

where $K^0$ is the partition coefficient at infinite dilution, $\phi_X^{\text{in}} \to 0$. Notice that the pure entropic exclusion term $(1/(1 - \phi_S^{\text{in}}))$ contributes a negative sign to the right-hand side, reflecting the free-energy cost of crowding. Therefore, a positive feedback (i.e., partitioning increasing with concentration) requires that the direct attractive interactions overcome this entropic penalty.

Exponentiating Eq. (S37) gives, to leading order,

$$K_{\text{eq}}(\phi_X^{\text{in}}) \equiv \frac{\phi_X^{\text{in}}}{\phi_X^{\text{out}}} \approx K^0 \exp\big(-\gamma_0^{\text{rep}}\,\phi_X^{\text{in}}\big), \tag{S38}$$

with

$$\gamma_0^{\text{rep}} \equiv NA = N\left[\frac{1}{1 - \phi_S^{\text{in}}} + (\chi_{XX} - \chi_{XW})\right]. \tag{S39}$$

To recover the exponential enhancement form used in the main text, we define the effective feedback coefficient $\gamma_0$ as the net coefficient after subtracting the repulsive contribution. Specifically, we write

$$\gamma_0 \equiv -NA = N\left[(\chi_{XW} - \chi_{XX}) - \frac{1}{1 - \phi_S^{\text{in}}}\right], \tag{S40}$$

so that the partition coefficient takes the canonical form

$$K_{\text{eq}}(\phi_X^{\text{in}}) = K^0 \exp\big(\gamma_0\phi_X^{\text{in}}\big). \tag{S41}$$

Positive feedback ($\gamma_0 > 0$) thus requires $(\chi_{XW} - \chi_{XX}) > 1/(1-\phi_S^{\rm in})$, i.e., the client–solvent interaction asymmetry must be sufficiently large, which is physically realized when client–client attractions ($\chi_{XX} < 0$) are strong.

If we retain the quadratic term from Eq. (S35), the equilibrium condition becomes

$$\ln \frac{K_{\rm eq}(\phi_X^{\rm in})}{K^0} = \gamma_0 \phi_X^{\rm in} - b_0 (\phi_X^{\rm in})^2 + \mathcal{O}\big((\phi_X^{\rm in})^3\big), \tag{S42}$$

with the saturation coefficient

$$b_0 = \frac{N}{2(1-\phi_S^{\rm in})^2} > 0. \tag{S43}$$

The negative sign before $b_0$ reflects the physical fact that the excluded-volume enhancement saturates as the client occupies an increasing fraction of the pore space and the entropy gain from further displacing solvent diminishes.

It is convenient to work with the dimensionless concentration $u$ normalised by the Michaelis constant $K_M$ of the client for the chaperone,

$$\phi_X^{\rm in} = v c^{\rm in} = v K_M\, u, \tag{S44}$$

where $v$ is the molecular volume of the client. Substituting into Eqs. (S41) and (S42), we define

$$\gamma \equiv \gamma_0\, v K_M = N v K_M \left[ (\chi_{XW} - \chi_{XX}) - \frac{1}{1-\phi_S^{\rm in}} \right], \tag{S45}$$

and

$$\beta \equiv b_0\, (v K_M)^2 = \frac{N v^2 K_M^2}{2(1-\phi_S^{\rm in})^2}. \tag{S46}$$

To linear order we recover the compact exponential form

$$K_{\rm eq}(u) = K^0 e^{\gamma u}. \tag{S47}$$

The parameter $\gamma$ (dimensionless, $\gamma \geq 0$) measures the strength of the positive thermodynamic feedback. $\gamma = 0$ recovers the constant partition coefficient of the linear theory. For $\gamma > 0$, the more client that accumulates inside the condensate, the higher the condensate's affinity for that client, a positive feedback loop that can destabilise the uniform clearance state.

To second order the partition coefficient reads

$$K_{\rm eq}(u) = K^0 \exp\Big[\gamma u - \beta u^2 + \mathcal{O}(u^3)\Big]. \tag{S48}$$

The parameter $\beta > 0$ is the saturation coefficient, which always acts to reduce the partition coefficient at high $u$. The pure exponential model, Eq. (S47), is recovered in the limit $\beta \to 0$, which is valid when $\beta u^2 \ll \gamma u$, i.e., $u \ll \gamma/\beta$. The leading-order phenomenology of bistability is controlled by $\gamma$, with $\beta$ providing small quantitative corrections in the physically relevant regime $\gamma^2 \gg 2\beta$.

#### Steady-state equation and saddle-node bifurcation

**Validity of the low-load approximation.** The linear extraction kinetics used below requires $u = c^{\rm in}/K_M \ll 1$. In the pure exponential model, the cusp is located at $u_c = 2/\gamma$. Hence, the analytical threshold $\mathrm{Da}_{0c} = e^{-2}$ is strictly rigorous in the limit $\gamma \gg 2$ (i.e., weak feedback relative to the Michaelis constant). For $\gamma \sim \mathcal{O}(1)$, the cusp concentration lies outside the linear regime; in that case, the full Michaelis–Menten saturation must be retained (see Sec.  and Fig. 1c for numerical phase diagrams). The following analysis therefore provides a benchmark asymptotics, which we later generalize numerically.

We now insert the concentration-dependent partition coefficient $K_{\rm eq}(u) = K^0 e^{\gamma u}$ into the single-client MNSR equation. The external supply is taken to be dilute ($u^{\rm out} \ll 1$) and the client load is low enough that chaperone saturation is negligible ($u \ll 1$, so that $1 + u \approx 1$). Extraction follows linear kinetics, $\mathcal{R}^{\rm cons} = V k_{\rm ext} c^{\rm in}$. Under these conditions the MNSR equation (S31) reduces to

$$k_{\rm mt} \left( u^{\rm out} - \frac{u}{K_{\rm eq}(u)} \right) = k_{\rm ext} u. \tag{S49}$$

It is natural to introduce two dimensionless parameters that combine the microscopic rates. The dimensionless external supply is

$$a \equiv K^0 u^{\text{out}}, \tag{S50}$$

which measures the equilibrium concentration that the client would have inside the condensate in the absence of extraction and feedback ($u_{\text{eq}} = K^0 u^{\text{out}} = a$ at $\gamma = 0$, $\mathrm{Da}_0 = 0$). The clearance strength (or reference Damköhler number based on the zero-concentration partition coefficient) is

$$\mathrm{Da}_0 \equiv \frac{K^0 k_{\text{ext}}}{k_{\text{mt}}}. \tag{S51}$$

$\mathrm{Da}_0$ quantifies the rate of active extraction relative to diffusive resupply, weighted by the baseline affinity. Inserting $K_{\text{eq}}(u) = K^0 e^{\gamma u}$ and the definitions of $a$ and $\mathrm{Da}_0$ into Eq. (S49) yields, after multiplying through by $K^0/k_{\text{mt}}$,

$$a = \Phi(u) \equiv u e^{-\gamma u} + \mathrm{Da}_0\, u. \tag{S52}$$

Equation (S52) is the central equation of this section. The steady states of the system are the solutions $u \geq 0$ of $a = \Phi(u)$. The left-hand side $a$ is the external control parameter (proportional to the client concentration in the reservoir), while the right-hand side $\Phi(u)$ encapsulates the combined effect of thermodynamic feedback (the term $ue^{-\gamma u}$) and active extraction (the term $\mathrm{Da}_0\, u$).

To determine the number and stability of steady states we examine the shape of $\Phi(u)$ for $u \geq 0$. We have $\Phi(0) = 0$ and the asymptotic behaviour $\Phi(u) \sim \mathrm{Da}_0\, u$ for $u \to \infty$, since the exponential term $ue^{-\gamma u}$ vanishes at large $u$. At large concentrations $\Phi(u)$ grows linearly with slope $\mathrm{Da}_0$. At the origin the derivative is $\Phi'(0) = 1 + \mathrm{Da}_0 > 1$, so the curve initially rises with a slope greater than unity.

The derivative of $\Phi(u)$ at arbitrary $u$ is

$$\Phi'(u) = (1 - \gamma u) e^{-\gamma u} + \mathrm{Da}_0. \tag{S53}$$

For $\Phi(u)$ to be non-monotonic, a necessary condition for bistability, $\Phi'(u)$ must become negative for some range of $u$. Since $\mathrm{Da}_0 \geq 0$ and $e^{-\gamma u} > 0$, the only way $\Phi'(u)$ can be negative is if $(1 - \gamma u) < 0$, i.e., $u > 1/\gamma$, and simultaneously $\mathrm{Da}_0 < (\gamma u - 1) e^{-\gamma u}$.

The extrema of $\Phi(u)$, where $\Phi'(u) = 0$, satisfy

$$\mathrm{Da}_0 = h(\gamma u), \qquad h(z) \equiv (z - 1) e^{-z}, \quad z \geq 1. \tag{S54}$$

The function $h(z)$ plays a central role. Its key properties are that $h(0) = -1$, the root $z = 1$ gives $h(1) = 0$, $h(z) > 0$ for $z > 1$, and $h(z)$ attains a unique maximum at $z = 2$ where $h(2) = e^{-2} \approx 0.1353$. For $z \to \infty$, $h(z) \to 0^+$. Because $\mathrm{Da}_0 \geq 0$, Eq. (S54) has no solutions for $\mathrm{Da}_0 > e^{-2}$, in which case $\Phi'(u) > 0$ everywhere and $\Phi(u)$ is strictly monotonically increasing. At $\mathrm{Da}_0 = e^{-2}$ there is exactly one degenerate solution where $\Phi'(u) = 0$ at $u = 2/\gamma$ and $\Phi(u)$ has an inflection point with horizontal tangent. For $0 < \mathrm{Da}_0 < e^{-2}$ there are two distinct solutions $u_1 < u_2$, giving $\Phi(u)$ a local maximum at $u_1$ and a local minimum at $u_2$, with the ordering $0 < u_1 < 1/\gamma < 2/\gamma < u_2 < \infty$. The values $u_1$ and $u_2$ are the two positive roots of $\mathrm{Da}_0 = (\gamma u - 1) e^{-\gamma u}$.

Linear stability analysis shows that a steady state $u^*$ is stable if $\Phi'(u^*) > 0$ and unstable if $\Phi'(u^*) < 0$. Therefore, when $0 < \mathrm{Da}_0 < e^{-2}$ and $\Phi(u)$ is non-monotonic, the three steady-state branches are a stable lower branch $u \in [0, u_1)$ (the clearance state $u_{\text{low}}$), an unstable middle branch $u \in (u_1, u_2)$ (the saddle $u_{\text{barr}}$), and a stable upper branch $u \in (u_2, \infty)$ (the storage state $u_{\text{high}}$).

The bistable region in the $(\mathrm{Da}_0, a)$ parameter plane is defined by the condition that three steady states coexist. This requires $0 < \mathrm{Da}_0 < e^{-2}$ and

$$a_{\min}(\mathrm{Da}_0) < a < a_{\max}(\mathrm{Da}_0), \tag{S55}$$

where

$$a_{\min}(\mathrm{Da}_0) \equiv \Phi(u_2(\mathrm{Da}_0)), \qquad a_{\max}(\mathrm{Da}_0) \equiv \Phi(u_1(\mathrm{Da}_0)). \tag{S56}$$

Although $u_1$ and $u_2$ are given implicitly, the boundaries can be expressed in a closed parametric form. Let $s = \gamma u$ be the rescaled concentration at an extremum. From $\Phi'(u) = 0$ we have $\mathrm{Da}_0 = (s - 1) e^{-s}$ and the value of $\Phi$ at the extremum is $\gamma \Phi(u) = s e^{-s} + \mathrm{Da}_0 s = s^2 e^{-s}$. Thus the boundaries of the bistable window are

$$a = \frac{1}{\gamma}\, s^2 e^{-s}, \qquad \mathrm{Da}_0 = (s - 1) e^{-s}, \qquad s \geq 1. \tag{S57}$$

The upper boundary $a_{\max}(\mathrm{Da}_0)$ corresponds to $s \in [1,2]$ and the lower boundary $a_{\min}(\mathrm{Da}_0)$ to $s \in [2,\infty)$. At $s=1$, $\mathrm{Da}_0 = 0$ and $a = 1/(\gamma e)$; at $s=2$, $\mathrm{Da}_0 = e^{-2}$ and $a = 4/(\gamma e^2)$; as $s \to \infty$, $\mathrm{Da}_0 \to 0^+$ and $a \to 0^+$. The two branches meet at the cusp point

$$(\mathrm{Da}_{0c}, a_c) = \left(e^{-2}, \frac{4}{\gamma e^2}\right). \tag{S58}$$

For $\mathrm{Da}_0 > e^{-2}$ no bistability exists regardless of $a$. For $\mathrm{Da}_0 < e^{-2}$ bistability exists for $a$ in a finite interval whose width $\Delta a = a_{\max} - a_{\min}$ grows as $(e^{-2} - \mathrm{Da}_0)^{3/2}$ near the cusp, the classic scaling of a saddle-node bifurcation.

A remarkable feature is that the threshold $\mathrm{Da}_{0c} = e^{-2}$ is universal within the exponential feedback class. It does not depend on the microscopic feedback strength $\gamma$, which only rescales the concentration and supply axes. Physically, $\mathrm{Da}_0$ can be expressed in terms of the fundamental dimensionless parameters as $\mathrm{Da}_0 = K^0\alpha^2\rho/3$, where $\alpha = R\sqrt{k_{\mathrm{ext}}/D^{\mathrm{in}}}$ is the Thiele modulus and $\rho = D^{\mathrm{in}}/D^{\mathrm{out}}$ is the diffusivity ratio. The threshold $\mathrm{Da}_0 = e^{-2}$ therefore translates into a condition on droplet size, extraction rate, and diffusion coefficients.

#### Corrections to the ideal threshold

The condition $\mathrm{Da}_0 < e^{-2}$ was derived under three idealisations: purely exponential feedback, linear extraction kinetics, and a spatially uniform intra-condensate concentration ($\alpha \ll 1$). In real condensates these assumptions are violated. While a strict mathematical factorization into a single effective Damköhler number is not generally valid for arbitrary parameter ranges, we can provide approximate heuristic corrections that illustrate the direction of each effect.

**Internal gradients.** When the Thiele modulus $\alpha = R\sqrt{k_{\mathrm{ext}}/D^{\mathrm{in}}}$ is not negligible, the client concentration becomes non-uniform inside the droplet. Solving the reaction–diffusion equation $D^{\mathrm{in}}\nabla^2 c = k_{\mathrm{ext}}c$ in spherical geometry gives the effectiveness factor

$$\eta(\alpha) = \frac{3}{\alpha}\left(\frac{1}{\tanh\alpha} - \frac{1}{\alpha}\right) \le 1. \tag{S59}$$

This modifies the steady-state balance to $a = ue^{-\gamma u} + \mathrm{Da}_0\,\eta(\alpha)\,u$, weakening the effective clearance.

**Michaelis–Menten saturation.** For finite client loads, the extraction term becomes $k_{\mathrm{ext}}\,u/(1+u)$. The steady-state equation reads $a = ue^{-\gamma u} + \mathrm{Da}_0\,u/(1+u)$. In contrast to a simple multiplicative rescaling, the exact bifurcation threshold must be obtained by solving $\Phi'(u) = 0$ and $\Phi''(u) = 0$ simultaneously. For example, for $\gamma = 1$, the full saturation model yields $\mathrm{Da}_{0c} = 32e^{-3} \approx 1.593$, whereas the linearized estimate $e^{-2}$ holds only for $\gamma \gg 2$ (where $u_c = 2/\gamma \ll 1$).

**Quadratic feedback saturation.** Retaining the quadratic term of the Flory–Huggins expansion gives $K_{\mathrm{eq}}(u) = K^0\exp[\gamma u - \beta u^2]$ with $\beta > 0$ defined microscopically in Eqs. (S45)–(S46). To leading order in $\beta$, the critical clearance shifts as

$$\mathrm{Da}_{0c} \approx e^{-2}\left(1 - \frac{4\beta}{\gamma^2}\right), \tag{S60}$$

which reduces the bistable window.

**Heuristic multiplicative form and its limitations.** If one formally combines the above leading-order corrections into a single effective number, one obtains a heuristic expression

$$\mathrm{Da}_{\mathrm{eff}}^{\mathrm{heur}} \approx \mathrm{Da}_0\,\eta(\alpha)\,\frac{1}{1+u_{\mathrm{char}}}\left(1 - \frac{4\beta}{\gamma^2}\right)^{-1}, \quad \text{with } u_{\mathrm{char}} = 2/\gamma. \tag{S61}$$

We emphasize that **this expression is not a rigorous mathematical result** for the full nonlinear system. It serves only as a qualitative guide to the direction of each correction. For quantitative predictions and for parameter sets where $\gamma \sim \mathcal{O}(1)$ or $\alpha \gtrsim 1$, the full steady-state equation must be solved numerically. The phase diagrams in Fig. 1(c)–(e) are generated from such exact numerical solutions, and they show that while the pristine $e^{-2}$ value is shifted, the saddle-node bifurcation topology and the associated scaling laws remain robust.

For a mixture of $m$ clients with feedback matrix $\Gamma_{ij} \ge 0$, under the dominant-mode approximation the collective coordinate $U = \sum_i v_i^{(1)} u_i$ associated with the largest eigenvalue $\lambda_{\max}$ follows a single-client equation with effective parameters $\tilde{a}$, $\tilde{\mathrm{Da}}_0$, $\tilde{u}_{\mathrm{char}}$, and $\tilde{\beta}$ defined analogously. The heuristic threshold then reads $\tilde{\mathrm{Da}}_{\mathrm{eff}}^{\mathrm{heur}} < e^{-2}$, with the same caveats regarding its approximate nature.

#### Effective potential and energy landscape

The deterministic dynamics of the single-client concentration $u$ can be written as

$$\tau_0 \frac{du}{dt} = a - \Phi(u) = -\frac{dU_{\rm eff}}{du}, \qquad U_{\rm eff}(u) \equiv \int_0^u \big[\Phi(s) - a\big]\, ds. \tag{S62}$$

Using $\Phi(u) = ue^{-\gamma u} + \mathrm{Da}_0\, u$, the integral evaluates to

$$U_{\rm eff}(u) = \frac{1}{\gamma^2}\Big[1 - (1+\gamma u)e^{-\gamma u}\Big] + \frac{\mathrm{Da}_0}{2}u^2 - au, \tag{S63}$$

with $U_{\rm eff}(0) = 0$. In the bistable regime ($0 < \mathrm{Da}_0 < e^{-2}$, $a_{\rm min} < a < a_{\rm max}$) this potential has two minima, the clearance state $u_{\rm low}$ and the storage state $u_{\rm high}$, separated by a maximum at $u_{\rm barr}$, precisely the fixed points analysed above. The forward and reverse barriers are $\Delta U_{\rm for} = U_{\rm eff}(u_{\rm barr}) - U_{\rm eff}(u_{\rm low})$ and $\Delta U_{\rm rev} = U_{\rm eff}(u_{\rm barr}) - U_{\rm eff}(u_{\rm high})$, and the free-energy bias between the two states is $\Delta U_{\rm bias} \equiv U_{\rm eff}(u_{\rm high}) - U_{\rm eff}(u_{\rm low}) = \Delta U_{\rm for} - \Delta U_{\rm rev}$. For multi-component condensates the same structure holds with the replacements $u \to U$, $\gamma \to \lambda_{\rm max}$, $\mathrm{Da}_0 \to \tilde{\mathrm{Da}}_0$, $a \to \tilde{a}$. This double-well landscape governs both deterministic relaxation and, in the presence of molecular noise, the stochastic transitions discussed in the following sections.

### FUNCTIONAL LIFETIME UNDER MOLECULAR NOISE

In the deterministic analysis of Sec. , a condensate placed in the clearance state will remain there forever in the absence of perturbations. Real condensates are finite, and intrinsic molecular noise can drive the system over the barrier even when the clearance state is deterministically stable, leading to a finite functional lifetime. We derive this lifetime from the stochastic MNSR model, first obtaining an effective Langevin dynamics, then applying Kramers' escape-rate theory. The result is a piecewise-defined expression for $\tau_{\rm func}$ covering the entire $(\mathrm{Da}_0, a)$ phase diagram, together with scaling laws near the saddle-node bifurcation and the non-monotonic ATP dependence.

#### From chemical master equation to effective Langevin dynamics

Let $n$ be the total number of client molecules inside the condensate and define the system size $\Omega \equiv VK_M$, where $V$ is the condensate volume and $K_M$ the Michaelis constant. The dimensionless concentration is $u = n/\Omega$. In the dilute-client, linear-extraction limit, the elementary events are diffusive entry, passive exit, and active extraction, with propensity functions

$$a_+(u) = \Omega\, k_{\rm mt}\, u^{\rm out}, \tag{S64}$$

$$a_-(u) = \Omega \left[\frac{k_{\rm mt}\, u}{K^0 e^{\gamma u}} + k_{\rm ext}\, u\right], \tag{S65}$$

where $u^{\rm out} = a/K^0$ and $a = K^0 u^{\rm out}$ is the dimensionless external supply introduced in the main text.

Expanding the chemical master equation to second order in $1/\Omega$ gives the Kramers–Moyal Fokker–Planck equation

$$\frac{\partial P(u,t)}{\partial t} = -\frac{\partial}{\partial u}\big[A(u)P\big] + \frac{1}{2\Omega}\frac{\partial^2}{\partial u^2}\big[B(u)P\big], \tag{S66}$$

where the drift and diffusion functions are

$$A(u) = \frac{a_+(u) - a_-(u)}{\Omega} = k_{\rm mt}u^{\rm out} - \frac{k_{\rm mt}u}{K^0 e^{\gamma u}} - k_{\rm ext}u = \frac{k_{\rm mt}}{K^0}\left[a - ue^{-\gamma u} - \frac{K^0 k_{\rm ext}}{k_{\rm mt}}u\right] = \frac{1}{\tau_0}\big[a - \Phi(u)\big], \tag{S67}$$

$$B(u) = \frac{a_+(u) + a_-(u)}{\Omega} = k_{\rm mt}u^{\rm out} + \frac{k_{\rm mt}u}{K^0 e^{\gamma u}} + k_{\rm ext}u, \tag{S68}$$

with $\tau_0 \equiv K^0/k_{\rm mt}$ and $\Phi(u) \equiv ue^{-\gamma u} + \mathrm{Da}_0 u$, where $\mathrm{Da}_0 = K^0 k_{\rm ext}/k_{\rm mt}$.

Linearising Eq. (S66) around a stable fixed point $u^*$, $a = \Phi(u^*)$, and approximating $B(u) \approx B(u^*)$, we obtain an Ornstein–Uhlenbeck process

$$\frac{d\delta u}{dt} = -\frac{\Phi'(u^*)}{\tau_0}\,\delta u + \sqrt{2\Lambda}\,\xi(t), \tag{S69}$$

where $\delta u = u - u^*$, $\xi(t)$ is Gaussian white noise, and the noise intensity is $\Lambda = B(u^*)/(2\Omega)$. The corresponding effective temperature $\Theta_{\text{eff}} \equiv \tau_0\Lambda$ then reads

$$\Theta_{\text{eff}} = \frac{\tau_0 B(u^*)}{2\Omega} = \frac{\tau_0}{2\Omega}\cdot 2k_{\text{mt}}u^{\text{out}} = \frac{a}{VK_M}, \tag{S70}$$

where we used the steady-state balance $a_+(u^*) = a_-(u^*)$, which gives $B(u^*) = 2k_{\text{mt}}u^{\text{out}} = 2k_{\text{mt}}a/K^0$, and $\tau_0 = K^0/k_{\text{mt}}$. Thus the effective temperature is fixed solely by the external supply $a$ and the number scale $VK_M$; it is independent of $\text{Da}_0$ and $\gamma$, and scales as

$$\Theta_{\text{eff}} \propto R^{-3}, \tag{S71}$$

where $R$ is the droplet radius.

#### Effective Boltzmann description and Ginzburg criterion

If $\Lambda$ is treated as constant, the steady-state solution of Eq. (S66) is

$$P_{\text{st}}(u) \propto \exp\left[-\frac{U_{\text{eff}}(u)}{\tau_0\Lambda}\right], \qquad U_{\text{eff}}(u) = \int_0^u [\Phi(s) - a]\,ds. \tag{S72}$$

This effective Boltzmann distribution is valid only when the noise intensity does not vary appreciably over the typical width of the potential well. The linear-noise approximation requires

$$|B'(u^*)|\sqrt{\frac{\tau_0\Lambda}{|U''_{\text{eff}}(u^*)|}} \ll B(u^*), \tag{S73}$$

which is satisfied for the exponential-feedback model whenever the well curvature is not too small. A more stringent Ginzburg criterion for the validity of Kramers' escape rate is that the effective temperature be small compared with both barrier heights:

$$\tau_0\Lambda \ll \Delta U_{\text{for}} \quad \text{and} \quad \tau_0\Lambda \ll \Delta U_{\text{rev}}. \tag{S74}$$

Equivalently, the typical fluctuation amplitude $\delta u_{\text{typ}} = \sqrt{\tau_0\Lambda/|U''_{\text{eff}}(u^*)|}$ must be much smaller than the distance between the fixed point and the saddle. Near a saddle-node bifurcation, both $\Delta U$ and $U''_{\text{eff}}(u^*)$ vanish, so the criterion (S74) eventually fails; there the effective Boltzmann description breaks down and the full Fokker–Planck equation must be used.

For typical condensate parameters $V \sim 1\,\mu\text{m}^3$, $K_M \sim 1\,\mu\text{M}$, and $a \sim 0.1$, the effective temperature is $\Theta_{\text{eff}} \sim 10^{-3}$. The barrier heights $\Delta U$ are of order 0.1–1 far from the bifurcation, so the Ginzburg criterion is safely satisfied. Only within a narrow critical region around $\text{Da}_{\text{eff}} = e^{-2}$ does the effective Boltzmann description lose quantitative accuracy. The key consequence for the functional lifetime is

$$\ln\tau_{\text{func}} \propto \frac{\Delta U_{\text{for}}}{\Theta_{\text{eff}}} \propto R^3\Delta U_{\text{for}}, \tag{S75}$$

so larger condensates are stabilized cubically with radius, making the upper size limit imposed by energetic cost even more significant.

#### Functional lifetime as a mean first-passage time

The functional state is identified with the low-concentration clearance minimum of $U_{\text{eff}}(u)$. We define the functional lifetime $\tau_{\text{func}}$ as the mean first-passage time from $u_{\text{low}}$ to the storage state $u_{\text{high}}$. Because $u_{\text{high}}$ lies beyond the barrier, the MFPT is dominated by the rare event of crossing the saddle at $u_{\text{barr}}$.

For an overdamped one-dimensional system with drift $-\frac{1}{\tau_0}U'_{\rm eff}(u)$ and diffusion constant $\Lambda$, the exact MFPT is

$$\tau_{\rm func}(u_{\rm low} \to u_{\rm high}) = \frac{1}{\Lambda}\int_{u_{\rm low}}^{u_{\rm high}} dy\, \exp\Big(\frac{U_{\rm eff}(y)}{\tau_0\Lambda}\Big)\int_{-\infty}^{y} dz\, \exp\Big(-\frac{U_{\rm eff}(z)}{\tau_0\Lambda}\Big). \tag{S76}$$

In the weak-noise limit $\tau_0\Lambda \ll \Delta U_{\rm for}$, Laplace's method yields the Kramers formula

$$\tau_{\rm func} = \frac{2\pi\,\tau_0}{\sqrt{\Phi'(u_{\rm low})\,|\Phi'(u_{\rm barr})|}}\,\exp\Big(\frac{\Delta U_{\rm for}}{\tau_0\Lambda}\Big) \equiv \tau_{\rm pref}\exp\Big(\frac{\Delta U_{\rm for}}{\tau_0\Lambda}\Big), \tag{S77}$$

where we used $U''_{\rm eff}(u) = \Phi'(u) = (1-\gamma u)e^{-\gamma u} + {\rm Da}_0$. The prefactor is

$$\tau_{\rm pref} \equiv \frac{2\pi\,\tau_0}{\sqrt{\Phi'(u_{\rm low})\,|\Phi'(u_{\rm barr})|}}, \tag{S78}$$

and the forward barrier height is

$$\Delta U_{\rm for} = \int_{u_{\rm low}}^{u_{\rm barr}} \big[\Phi(s) - a\big]\,ds. \tag{S79}$$

Equation (S77) is the central result for the functional lifetime in the bistable regime.

#### Lifetime in the complete $({\rm Da}_0, a)$ phase diagram

The stability landscape is fully determined by $\mathrm{Da}_0$ and $a$ (Sec. ). We can assign a functional lifetime to each point in this diagram, yielding a piecewise-defined function.

In the monostable clearance regime ($\mathrm{Da}_0 \geq e^{-2}$, or $\mathrm{Da}_0 < e^{-2}$ with $a \leq a_{\rm min}(\mathrm{Da}_0)$), no storage state exists and the mean first-passage time to $u \to \infty$ is infinite:

$$\tau_{\rm func} = \infty. \tag{S80}$$

In the monostable storage regime ($\mathrm{Da}_0 < e^{-2}$ with $a \geq a_{\rm max}(\mathrm{Da}_0)$), the clearance minimum has disappeared, so

$$\tau_{\rm func} = 0. \tag{S81}$$

In the bistable regime ($0 < \mathrm{Da}_0 < e^{-2}$ and $a_{\rm min}(\mathrm{Da}_0) < a < a_{\rm max}(\mathrm{Da}_0)$), the double-well potential is present and the functional lifetime is given by the Kramers formula (S77),

$$\tau_{\rm func}({\rm Da}_0, a) = \tau_{\rm pref}({\rm Da}_0, a)\,\exp\Big(\frac{\Delta U_{\rm for}({\rm Da}_0, a)}{\tau_0\Lambda}\Big). \tag{S82}$$

Within the bistable regime, $\tau_{\rm func}$ varies continuously from infinity on the lower boundary $a = a_{\rm min}$ (where $u_{\rm low}$ merges with the saddle) to zero on the upper boundary $a = a_{\rm max}$ (where the saddle merges with $u_{\rm high}$). The exponential dependence on the barrier height implies that the lifetime changes by many orders of magnitude across the narrow bistable window.

#### Asymptotic scaling laws for the barrier and the lifetime

Although the fixed points $u_{\rm low}$, $u_{\rm barr}$, and $u_{\rm high}$ are roots of $a = ue^{-\gamma u} + \mathrm{Da}_0\,u$ and cannot be expressed in elementary functions, their scaling near the boundaries of the bistable region can be obtained analytically.

Near the cusp $(\mathrm{Da}_{0c}, a_c) = (e^{-2}, 4/(\gamma e^2))$, we set $\mathrm{Da}_0 = e^{-2} - \varepsilon$ with $0 < \varepsilon \ll 1$. The shifted control parameter $B = \gamma(a - a_c) + 2\varepsilon$ governs the shape of the potential. The symmetric double-well configuration, in which the two minima have equal depth and the barrier is exactly halfway, corresponds to $B = 0$, i.e.

$$a = a_{\rm sym}(\varepsilon) = a_c - \frac{2\varepsilon}{\gamma}. \tag{S83}$$

Along this path the cubic normal form $B = \frac{e^{-2}}{6}x^3 - \varepsilon x$ with $x = \gamma u - 2$ yields three steady states: $x = 0$ (saddle) and $x = \pm e\sqrt{2\varepsilon}$ (the two symmetric minima). Inserting these into the effective potential $U_{\text{eff}}$ and performing the integral gives the barrier height

$$\Delta U_{\text{for}} = \frac{3e^2}{2\gamma^2}\,\varepsilon^2 = \frac{3e^2}{2\gamma^2}\,(e^{-2} - \text{Da}_0)^2. \tag{S84}$$

The prefactor $\tau_{\text{pref}}$ remains finite at $\text{Da}_0 = e^{-2}$, so the functional lifetime scales as

$$\tau_{\text{func}} \sim \exp\Big(\frac{3e^2}{2\gamma^2\tau_0\Lambda}\,(e^{-2} - \text{Da}_0)^2\Big). \tag{S85}$$

Using $\tau_0\Lambda \propto R^{-3}$ from Eq. (S71), the exponent contains the factor $R^3\,(e^{-2} - \text{Da}_0)^2$. Thus larger droplets are exponentially more stable against noise-induced failure.

When $a$ approaches $a_{\min}(\text{Da}_0)$ from above, the clearance minimum $u_{\text{low}}$ and the saddle $u_{\text{barr}}$ merge in a saddle-node bifurcation. Let $\delta a = a - a_{\min} \ll 1$. Standard bifurcation theory gives

$$u_{\text{barr}} - u_{\text{low}} \propto \delta a^{1/2}, \qquad \Phi'(u_{\text{low}}) \propto \delta a^{1/2}, \qquad \Delta U_{\text{for}} \propto \delta a^{3/2}. \tag{S86}$$

Substituting these into the Kramers formula (S77) yields

$$\tau_{\text{func}} \sim \delta a^{-1/4}\exp\Big(-\frac{c\,\delta a^{3/2}}{\tau_0\Lambda}\Big) \to 0 \quad \text{as } \delta a \to 0^+. \tag{S87}$$

Near the upper boundary $a \to a_{\max}(\text{Da}_0)$, the saddle and the storage minimum merge, while the clearance minimum remains well separated. The forward barrier $\Delta U_{\text{for}}$ scales as $(a_{\max} - a)^{3/2}$ and the prefactor diverges as $(a_{\max} - a)^{-1/4}$, leading to

$$\tau_{\text{func}} \sim (a_{\max} - a)^{-1/4}\exp\Big(-\frac{c'\,(a_{\max} - a)^{3/2}}{\tau_0\Lambda}\Big) \to 0 \quad \text{as } a \nearrow a_{\max}. \tag{S88}$$

The same scaling applies if the upper boundary is crossed by varying $\text{Da}_0$ while keeping $a$ fixed, with the distance parameter replaced by $\text{Da}_{0\,\max}(a) - \text{Da}_0$.

#### Non-monotonic ATP dependence of stability

ATP concentration $c_T$ influences the clearance strength $\text{Da}_0$ through two competing mechanisms: it activates chaperones so that $k_{\text{ext}}(c_T)$ increases, and it acts as a hydrotrope so that $K^0(c_T)$ decreases. The net effect is that $\text{Da}_0(c_T)$ is a unimodal function of $c_T$, with a single maximum $\text{Da}_{0\,\max}$ at an optimal ATP concentration $c_T^*$.

Inserting this $\text{Da}_0(c_T)$ dependence into the piecewise expression for $\tau_{\text{func}}$ reveals three qualitatively distinct ATP regimes. At low ATP, where $\text{Da}_0 \ll e^{-2}$, the system lies deep in the bistable regime and the barrier $\Delta U_{\text{for}}$ is large, of order $\gamma^{-2}$; the clearance state is kinetically stable. At intermediate ATP, where $\text{Da}_0 \lesssim e^{-2}$, the barrier collapses quadratically according to Eq. (S84), and $\tau_{\text{func}}$ drops exponentially. This is the window of maximal vulnerability, because a modest amount of ATP destabilises the clearance state, making it highly susceptible to noise, without eliminating the storage state. At high ATP, where $\text{Da}_0 \geq e^{-2}$, the storage state disappears entirely and the condensate enters the monostable functional regime with $\tau_{\text{func}} = \infty$.

This analysis yields a counterintuitive prediction: the onset of condensate dysfunction under metabolic stress may occur before ATP is completely depleted. Partial energy deprivation that places $\text{Da}_0$ just below $e^{-2}$ creates a window of maximal vulnerability. Complete ATP exhaustion, paradoxically, would restore a long-lived, though fragile, clearance state. This non-monotonic behaviour is a direct consequence of the saddle-node bifurcation and the unimodal nature of $\text{Da}_0(c_T)$.

#### Volume-dependent stability

Because the effective temperature scales as $\Theta_{\text{eff}} = \tau_0\Lambda \propto R^{-3}$, the functional lifetime depends exponentially on droplet radius as

$$\ln \tau_{\text{func}} \propto \frac{R^3\,\Delta U_{\text{for}}}{\text{const}}. \tag{S89}$$

Smaller condensates are exponentially more vulnerable to noise-induced functional loss. This provides a physical rationale for the observed size regulation of many cellular condensates: there exists a minimal droplet size below which the clearance state is no longer kinetically viable, setting a lower bound on the size of functional organelles. It also predicts that pathological aggregation should nucleate preferentially in smaller droplets, where the noise-driven escape from the functional state is fastest.

### ENERGETIC COST OF RECOVERY AND HYSTERESIS

The bistable landscape derived in Sec. implies that a condensate, once driven into the pathological storage state, cannot spontaneously return to the functional clearance state without external energy input. This section quantifies the minimal energetic cost of active recovery and the non-equilibrium work dissipated during cyclic stress-recovery protocols.

#### Reverse barrier and free-energy bias

In the bistable regime $0 < \mathrm{Da}_0 < e^{-2}$ and $a_{\min} < a < a_{\max}$, the effective potential $U_{\mathrm{eff}}(u)$ has two minima, $u_{\mathrm{low}}$ (clearance) and $u_{\mathrm{high}}$ (storage), separated by a maximum at $u_{\mathrm{barr}}$. The forward and reverse barriers and the free-energy bias are

$$\Delta U_{\mathrm{for}} = U_{\mathrm{eff}}(u_{\mathrm{barr}}) - U_{\mathrm{eff}}(u_{\mathrm{low}}) > 0, \tag{S90}$$

$$\Delta U_{\mathrm{rev}} = U_{\mathrm{eff}}(u_{\mathrm{barr}}) - U_{\mathrm{eff}}(u_{\mathrm{high}}) > 0, \tag{S91}$$

$$\Delta U_{\mathrm{bias}} \equiv U_{\mathrm{eff}}(u_{\mathrm{high}}) - U_{\mathrm{eff}}(u_{\mathrm{low}}) = \Delta U_{\mathrm{for}} - \Delta U_{\mathrm{rev}}. \tag{S92}$$

When $\Delta U_{\mathrm{bias}} < 0$, the storage state is the global free-energy minimum. Using the explicit form of $U_{\mathrm{eff}}$, the bias can be written as

$$\Delta U_{\mathrm{bias}} = \int_{u_{\mathrm{low}}}^{u_{\mathrm{high}}} \big[\Phi(s) - a\big]\, ds, \tag{S93}$$

and in the limit $\mathrm{Da}_0 \ll e^{-2}$, an asymptotic expansion gives

$$\Delta U_{\mathrm{bias}} \simeq -\frac{2}{\gamma^2}\Big[e^{-2} - \mathrm{Da}_0 + \mathrm{Da}_0 \ln\frac{\mathrm{Da}_0}{e^{-2}}\Big] < 0. \tag{S94}$$

The bias is negative and grows in magnitude as $\mathrm{Da}_0$ decreases; as $\mathrm{Da}_0 \nearrow e^{-2}$, $\Delta U_{\mathrm{bias}} \to 0$.

#### Minimal work for resetting: fluctuation-theorem bound

In practice, cells transiently upregulate chaperone activity, increasing $\mathrm{Da}_0$ above $e^{-2}$ to eliminate the storage state and drive the system into the clearance state. This active resetting protocol consumes ATP and performs work. The minimal work required can be bounded by non-equilibrium fluctuation theorems.

For a protocol in which $\mathrm{Da}_0(t)$ is temporarily raised above $e^{-2}$, the Crooks fluctuation theorem for the overdamped Langevin dynamics gives

$$\frac{P_{\mathrm{for}}(W)}{P_{\mathrm{rev}}(-W)} = \exp\Big(\frac{W - \Delta F}{\tau_0 \Lambda}\Big), \tag{S95}$$

where $\Delta F = U_{\mathrm{eff}}(u_{\mathrm{high}}) - U_{\mathrm{eff}}(u_{\mathrm{low}}) = \Delta U_{\mathrm{bias}}$. Jensen's inequality then implies

$$\langle W_{\mathrm{reset}}\rangle \geq -\Delta F = -\Delta U_{\mathrm{bias}} > 0. \tag{S96}$$

Thus the minimal work is exactly the magnitude of the free-energy bias. Substituting the asymptotic expression for the bias gives, for $\mathrm{Da}_0 \ll e^{-2}$,

$$\langle W_{\mathrm{reset}}\rangle \gtrsim \frac{2}{\gamma^2}\Big[e^{-2} - \mathrm{Da}_0 + \mathrm{Da}_0 \ln\frac{\mathrm{Da}_0}{e^{-2}}\Big]. \tag{S97}$$

As $\mathrm{Da}_0 \to 0$, this function approaches the finite constant $2e^{-2}/\gamma^2$, with a subleading correction $\propto \mathrm{Da}_0 \ln(\mathrm{Da}_0/e^{-2})$. Thus the resetting cost does not diverge in the weak-chaperone limit; it saturates at a value set by the cusp position.

**Hysteresis loop in a quasi-static stress-recovery cycle**

During a stress episode, the external supply $a$ of misfolded clients rises slowly and then returns to basal levels. If the variation of $a(t)$ is slow compared to $\tau_0$, the system tracks the stable branches of the bifurcation diagram quasi-statically, executing an irreversible hysteresis loop.

Starting from $a_0 < a_{\min}$, the system follows the clearance branch $u_{\text{low}}(a)$ until $a$ reaches $a_{\max}$, where it jumps to the storage branch $u_{\text{high}}(a_{\max})$. As $a$ decreases, it remains trapped on the storage branch until $a$ drops below $a_{\min}$, at which point it jumps back to $u_{\text{low}}(a_{\min})$. The energy dissipated in the two jumps is

$$\Delta E_{\uparrow} = U_{\text{eff}}\big(u_{\text{high}}(a_{\max})\big) - U_{\text{eff}}\big(u_{\text{low}}(a_{\max})\big), \tag{S98}$$

$$\Delta E_{\downarrow} = U_{\text{eff}}\big(u_{\text{low}}(a_{\min})\big) - U_{\text{eff}}\big(u_{\text{high}}(a_{\min})\big), \tag{S99}$$

and the total work dissipated per full cycle is

$$W_{\text{hyst}} = \Delta E_{\uparrow} + \Delta E_{\downarrow}. \tag{S100}$$

To extract the leading scaling behaviour, set $\text{Da}_0 = e^{-2} - \varepsilon$ with $0 < \varepsilon \ll 1$, and introduce the scaled variable $x = \gamma u - 2$. Expanding $\Phi(u)$ around $u_c = 2/\gamma$ gives the normal form

$$\gamma\big(a - a_c\big) = \frac{e^{-2}}{6}x^3 - \varepsilon x - 2\varepsilon + \mathcal{O}(\varepsilon x, x^4), \tag{S101}$$

with $a_c = 4/(\gamma e^2)$. Defining the shifted control parameter

$$B \equiv \gamma(a - a_c) + 2\varepsilon = \frac{e^{-2}}{6}x^3 - \varepsilon x, \tag{S102}$$

the saddle-node boundaries occur at $x = \pm\delta$ with $\delta = e\sqrt{2\varepsilon}$, corresponding to $B_{\max} = -B_{\min} = \frac{2\sqrt{2}\,e}{3}\,\varepsilon^{3/2}$.

In the quasi-static limit, the total dissipated work equals the area enclosed by the hysteresis loop in the $(a, u)$ plane:

$$W_{\text{hyst}} = \oint u\, da = \frac{1}{\gamma^2}\oint x\, dB = \frac{2}{\gamma^2}\int_0^{B_{\max}} \big[x_+(B) - x_-(B)\big]\, dB = \frac{9e^2}{\gamma^2}\,\varepsilon^2 + \mathcal{O}(\varepsilon^{5/2}), \tag{S103}$$

where $x_+(B) > x_-(B)$ are the two outer roots of $B = \frac{e^{-2}}{6}x^3 - \varepsilon x$. Thus the hysteresis area scales as

$$W_{\text{hyst}} \simeq \frac{9e^2}{\gamma^2}\,(e^{-2} - \text{Da}_0)^2. \tag{S104}$$

This quadratic scaling shows that the energy cost grows quadratically with the distance from the critical point, so a modest decline in chaperone activity dramatically inflates the metabolic price of restoring condensate function.

To convert to physical units, we use the effective temperature $\Theta_{\text{eff}} = \tau_0\Lambda = a/(VK_M)$ derived in Appendix . The physical free energy corresponding to the dimensionless potential $U_{\text{eff}}$ is

$$\mathcal{F} = k_{\text{B}}T \cdot \frac{U_{\text{eff}}}{\tau_0\Lambda} = k_{\text{B}}T \cdot \frac{VK_M}{a}\,U_{\text{eff}}. \tag{S105}$$

Hence the dimensionless work $W_{\text{hyst}}$ corresponds to a physical dissipation

$$W_{\text{hyst}}^{(\text{physical})} = k_{\text{B}}T \cdot \frac{VK_M}{a}\,W_{\text{hyst}}, \tag{S106}$$

which scales as $R^3$, not linearly. The equivalent number of ATP molecules consumed is $W_{\text{hyst}}^{(\text{physical})}/\Delta G_{\text{ATP}}$. This cubic size scaling provides a physical rationale for the upper limit on condensate size: larger droplets, though exponentially more stable against noise, require proportionally more metabolic energy to be cycled through a stress-recovery event.

#### Comparing passive escape and active resetting

The passive escape time from the storage state back to the clearance state is given by the Kramers formula with the reverse barrier $\Delta U_{\rm rev}$:

$$\tau_{\rm rev} = \tau_{\rm pref,rev} \exp\Big(\frac{\Delta U_{\rm rev}}{\tau_0 \Lambda}\Big). \tag{S107}$$

In the asymmetric case where the storage state is favoured, $\Delta U_{\rm bias} < 0$, we have $\Delta U_{\rm rev} = \Delta U_{\rm for} - \Delta U_{\rm bias} > \Delta U_{\rm for}$, making passive recovery exponentially slower than contamination. Active resetting, by contrast, imposes a one-time cost $\langle W_{\rm reset}\rangle \geq -\Delta U_{\rm bias}$ but achieves recovery on a fast, tunable timescale. The biological choice between these strategies depends on $\mathrm{Da}_0$ and the condensate volume $V$: for small condensates passive escape may be fast enough, while for large condensates active resetting is the only viable mechanism despite its large ATP cost.

#### Consequences for cellular energy budget and ageing

In aged or stressed cells, declining chaperone activity $k_{\rm ext}$ lowers $\mathrm{Da}_0$, causing the resetting work to increase roughly as $1/\mathrm{Da}_0$ and the hysteresis loss to increase quadratically. Disease-linked mutations that strengthen homotypic interactions increase $\gamma$, reducing the barrier height but inflating the recovery cost via the $\gamma^{-2}$ factor in Eqs. (S97) and (S104). Frequent stress-recovery cycles impose a non-negligible ATP demand, potentially contributing to the selective vulnerability of neurons in neurodegenerative disease.

### CRITICAL FLUCTUATIONS AND EARLY WARNING SIGNALS

The saddle-node bifurcation organising bistability is a critical point. Approaching it from the monostable side produces critical slowing down and diverging fluctuations, which serve as early warning signals. We derive the linearised stochastic dynamics and scaling laws for three distinct paths.

#### Linearised Ornstein–Uhlenbeck dynamics

In the monostable clearance regime, expanding around the stable fixed point $u^*$ gives

$$a - \Phi(u) = -\Phi'(u^*)\,\delta u + \mathcal{O}(\delta u^2), \tag{S108}$$

so the Langevin equation becomes

$$\frac{d\delta u}{dt} = -\frac{\Phi'(u^*)}{\tau_0}\,\delta u + \sqrt{2\Lambda}\,\xi(t), \tag{S109}$$

where $\delta u = u - u^*$. This Ornstein–Uhlenbeck process has steady-state variance

$$\langle (\delta u)^2 \rangle = \frac{\tau_0 \Lambda}{\Phi'(u^*)} = \frac{\Theta_{\rm eff}}{\Phi'(u^*)} \propto \frac{1}{\Phi'(u^*)}, \tag{S110}$$

and relaxation time

$$\tau_{\rm relax} = \frac{\tau_0}{\Phi'(u^*)} \propto \frac{1}{\Phi'(u^*)}, \tag{S111}$$

where we used the effective temperature $\Theta_{\rm eff} = \tau_0 \Lambda$ from Appendix . Thus both observables are controlled solely by the local slope $\Phi'(u^*)$.

**Divergence upon approaching the bifurcation**

As $\Phi'(u^*) \to 0$, both variance and relaxation time diverge, with path-dependent exponents.

*Cusp, symmetric* $\mathrm{Da}_0$*-direction.* Setting $\mathrm{Da}_0 = e^{-2} - \varepsilon$ and tuning the supply to keep the fixed point at the inflection point $u^* = 2/\gamma$,

$$a = a_c - \frac{2(e^{-2} - \mathrm{Da}_0)}{\gamma}, \tag{S112}$$

the slope becomes

$$\Phi'(u^*) = \frac{|\varepsilon|}{\gamma} \propto |\mathrm{Da}_0 - e^{-2}|. \tag{S113}$$

Hence

$$\langle(\delta u)^2\rangle \propto |\mathrm{Da}_0 - e^{-2}|^{-1}, \qquad \tau_{\mathrm{relax}} \propto |\mathrm{Da}_0 - e^{-2}|^{-1}. \tag{S114}$$

*Cusp, a-direction at fixed* $\mathrm{Da}_0 = e^{-2}$. With $\delta a = a - a_c$, the normal form expansion gives

$$\Phi(u) - a_c \propto \frac{e^{-2}}{6\gamma}[\gamma(u - u_c)]^3, \tag{S115}$$

so the fixed point obeys $\gamma(u^* - u_c) \sim \delta a^{1/3}$, and

$$\Phi'(u^*) \propto |\delta a|^{2/3}. \tag{S116}$$

Therefore

$$\langle(\delta u)^2\rangle \propto |a - a_c|^{-2/3}, \qquad \tau_{\mathrm{relax}} \propto |a - a_c|^{-2/3}. \tag{S117}$$

*Saddle-node boundaries.* Approaching any saddle-node line from the monostable side, the fixed point merges with the saddle, giving a quadratic tangency. With distance $\delta m$ to the boundary,

$$\Phi'(u^*) \propto \sqrt{\delta m}. \tag{S118}$$

Thus

$$\langle(\delta u)^2\rangle \propto \delta m^{-1/2}, \qquad \tau_{\mathrm{relax}} \propto \delta m^{-1/2}. \tag{S119}$$

This applies to $a \to a_{\max}$ or $a \to a_{\min}$, and to $\mathrm{Da}_0$ variations across the boundaries, with $\delta m$ the corresponding distance.

In summary, the critical exponents are $-1$ for the symmetric cusp approach, $-2/3$ for the cusp along $a$, and $-1/2$ for generic saddle-node boundaries.
